# Engineering the highly productive cyanobacterium *Synechococcus* sp. PCC 11901

**DOI:** 10.1101/2023.08.04.552009

**Authors:** Angelo J. Victoria, Tiago Toscano Selão, José Ángel Moreno-Cabezuelo, Lauren A. Mills, Grant A. R. Gale, David J. Lea-Smith, Alistair J. McCormick

**Affiliations:** Institute of Molecular Plant Sciences, School of Biological Sciences, University of Edinburgh, EH9 3BF, UK; Centre for Engineering Biology, School of Biological Sciences, University of Edinburgh, EH9 3BF, UK; Department of Chemical and Environmental Engineering, University of Nottingham, Nottingham NG7 2RD, UK; School of Biological Sciences, University of East Anglia, Norwich Research Park, Norwich, NR4 7TJ, UK

## Abstract

*Synechococcus* sp. PCC 11901 (PCC 11901) is a fast-growing marine cyanobacterial strain that has a capacity for sustained biomass accumulation to very high cell densities, comparable to that achieved by commercially relevant heterotrophic organisms. However, genetic tools to engineer PCC 11901 for biotechnology applications are limited. Here we describe a suite of tools based on the CyanoGate MoClo system to unlock the engineering potential of PCC 11901. First, we characterised neutral sites suitable for stable genomic integration that do not affect growth even at high cell densities. Second, we tested a suite of constitutive promoters, terminators, and inducible promoters including a 2,4-diacetylphloroglucinol (DAPG)-inducible PhlF repressor system, which has not previously been demonstrated in cyanobacteria and showed tight regulation and a 228-fold dynamic range of induction. Lastly, we developed a DAPG-inducible dCas9-based CRISPR interference (CRISPRi) system and a modular method to generate markerless mutants using CRISPR-Cas12a. Based on our findings, PCC 11901 is highly responsive to CRISPRi-based repression and showed high efficiencies for single insertion (31-81%) and multiplex double insertion (25%) genome editing with Cas12a. We envision that these tools will lay the foundations for the adoption of PCC 11901 as a robust model strain for engineering biology and green biotechnology.

**On sentence summary:** Genetic parts were characterised in *Synechococcus* sp. PCC 11901, including a tightly regulated inducible promoter system, efficient CRISPRi and a novel markerless Cas12a genome editing approach.

## INTRODUCTION

Climate change has necessitated a global shift towards more sustainable production practices and the building of a bioeconomy centred on Net Zero Carbon policies (*Net Zero Strategy*, 2021). Cyanobacteria are an attractive alternative to heterotrophic microbial bioproduction chassis, such as *Escherichia coli* and yeast, due to their capacity for biology-based carbon capture and utilisation and potential for the production of a wide array of useful chemicals (Zhang et al., 2017; Daneshvar et al., 2022). Unicellular model cyanobacterial strains, such as *Synechocystis* sp. PCC 6803 (hereafter PCC 6803) and *Synechococcus elongatus* PCC 7942, have been investigated as biorefineries for bulk commodity products such as biofuels (Kopka et al., 2017), bioplastics (Khetkorn et al., 2016), natural colourants (Puzorjov et al., 2022) and terpenoids (Rautela and Kumar, 2022), while filamentous strains such as *Arthrospira platensis* have been developed for the production of food and high-value therapeutic antibodies (Jester et al., 2022; Saveria et al., 2022). Nevertheless, slow growth rates and low biomass productivity remain key bottlenecks that limit the economic competitiveness and commercial expansion of cyanobacterial biotechnology (Lea-Smith et al., 2021).

Several fast-growing cyanobacterial strains have been reported that show strong potential for overcoming yield challenges (for recent review see Selão, 2022), including fresh water strains *Synechococcus elongatus* UTEX 2973 (hereafter UTEX 2973) (Yu et al., 2015), *Synechococcus elongatus* PCC 11801 (hereafter PCC 11801) (Jaiswal et al., 2018), *Synechococcus elongatus* PCC 11802 (Jaiswal et al., 2020), and marine strains *Synechococcus* sp. PCC 7002 (hereafter PCC 7002) (Batterton and Van Baalen, 1971) and *Synechococcus* sp. PCC 11901 (hereafter PCC 11901) (Włodarczyk et al., 2020). Marine cyanobacteria are of particular interest as they can utilise sea/brackish water, circumventing the need for freshwater resources (Hitchcock et al., 2020). In contrast to other fast-growing strains, PCC 11901 has an additional capacity for sustained growth to high densities (up to 30 g/L dry cell weight, DCW), similar to the maximum biomass accumulation observed for batch-cultured *E. coli* cultures (24.7 g/L DCW) (Soini et al., 2008). Furthermore, PCC 11901 can tolerate high light intensities (>900 μmol photons m^−2^ s^−1^), temperatures (up to 43°C), and salinities over 2-fold higher than sea water (Cho et al., 2023). PCC 11901 is amenable to natural transformation and its genome is fully-sequenced (Włodarczyk et al., 2020). PCC 11901 has also been engineered to produce free fatty acids yielding >6 mM (1.5 g L^−1^), which is comparable to that achieved by similarly engineered heterotrophic organisms (Xiao et al., 2018). The recent isolation of a cobalamin-independent strain of PCC 11901 (*Synechococcus* sp. UTEX 3154) that does not require the addition of vitamin B12 will help reduce scale-up costs (Mills et al., 2022). Together, these qualities make PCC 11901 an attractive model species for fundamental research and biotechnology applications.

In recent years, standardised molecular tools have been developed to help progress synthetic/engineering biology in cyanobacteria, such as the CyanoGate Modular Cloning system (Vasudevan et al., 2019) and the Start-Stop Assembly method (Taylor et al., 2021). Additional advances include the development of self-replicating vectors for extrachromosomal expression (Taton et al., 2014; Opel et al., 2022), recombineering (Jones et al., 2021), genetic circuits (Taton et al., 2017; Zhang et al., 2022), CRISPR interference (CRISPRi) (Yao et al., 2020), and CRISPR-based gene editing tools to reprogram metabolism (Baldanta et al., 2022; Cengic et al., 2022; Wang et al., 2023). Strategies for biocontainment of mutant cyanobacterial strains have also been developed, for example, using inducible kill-switches (Zhou et al., 2019). Nevertheless, such tools are often not readily transferrable between different species, and to date very few engineering approaches have been characterised in PCC 11901. Notably, Mills et al. (2022) recently reported that PCC 11901 was not compatible with the negative selection markers *sacB* and *codA* used for generating markerless mutants (i.e. a genome-modified mutant that lacks a selective antibiotic resistance cassette) that have key advantages for biotechnology applications (Lea-Smith et al., 2016).

In this study we investigated a broad set of available genetic parts and developed several novel tools for engineering PCC 11901. We tested five putative neutral integration sites to serve as loci for introducing heterologous DNA and explored the amenability of PCC 11901 to transconjugation. We then characterised the functionality of a suite of known and new genetic parts, including the 2,4-diacetylphloroglucinol (DAPG)-inducible PhlF repressor system. Building on this, we developed an inducible CRISPRi gene repression system and a novel hybrid plasmid approach to successfully generate markerless mutants using CRISPR-Cas12a. Together these tools should fast-track the further development of PCC 11901 as a commercially viable chassis strain for cyanobacterial biotechnology.

## RESULTS AND DISCUSSION

### Identification of robust neutral sites in PCC 11901 suitable for high-density growth

PCC 11901 is naturally transformable but exhibits partial resistance to kanamycin (Włodarczyk et al., 2020). We first tested the susceptibility of PCC 11901 to five common selective antibiotics used to generate marked mutants (**Figure 1A**). We observed that PCC 11901 exhibited partial resistance to kanamycin at 50 µg mL^−1^ and gentamicin at 10 µg mL^−1^. In contrast, PC11901 was fully susceptible to spectinomycin, erythromycin, and chloramphenicol at all concentrations tested. Previous work has reported that many filamentous and unicellular species have innate resistances to specific antibiotics, including aminoglycosides such as kanamycin and gentamicin (Dias et al., 2015). Aminoglycoside resistance has been attributed to native redox-active compounds such as glutathione in *Synechocystis* sp. PCC 6803 (Cameron and Pakrasi, 2011), and the presence of native resistance genes such as the kanamycin aminoglycoside acetyltransferase homolog recently described in *Arthrospira platensis* (NIES39_D01030, GenBank ID: BAI89523.1) (Jester et al., 2022; Shiraishi and Nishida, 2022), which is a member of the Gcn5-related N-acetyltransferase (GNAT) superfamily (Favrot et al., 2016). A BLAST analysis of the PCC 11901 genome yielded 13 genes belonging to the GNAT family. The closest homolog to BAI89523.1 (FEK30_08270, GenBank ID: QCS49435.1) shared a 33% peptide sequence identity with similar conserved domains and a coenzyme A binding pocket motif, and thus may account for the partial resistance observed (**Supplementary Figure S1**).

**Figure 1.**
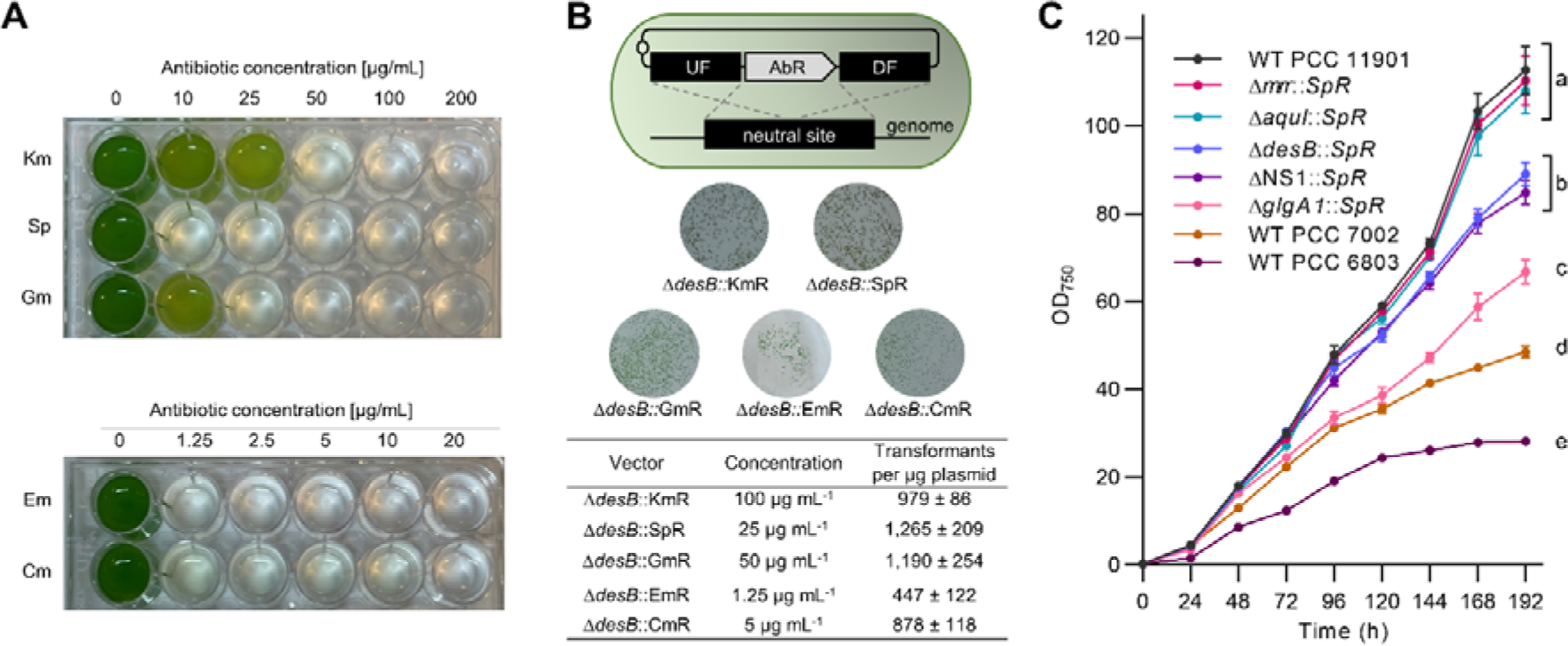
Antibiotic susceptibility and characterisation of putative neutral integration sites in PCC 11901. **(A)** Susceptibility of wild-type PCC 11901 to increasing concentrations of common antibiotics. PCC 11901 wild-type cultures were inoculated at OD_750_ = 0.2 and grown in MAD medium as described in the Materials and Methods for 48 h. **(B)** The diagram illustrates the transformation strategy used to introduce antibiotic resistance (AbR) cassettes into each putative neutral site via homologous recombination. An integrative pUC19 plasmid vector was assembled using the CyanoGate MoClo system (Vasudevan et al., 2019) and 1 µg of each plasmid was transformed into wild-type (WT) PCC 11901 (see **Supplementary Table S2** for plasmid vectors). Colony images and numbers of mutants transformed with different AbR cassettes integrated into the *desB* neutral site. Colony counts were estimated by dividing the plate into nine sectors and taking the average colony counts of three sectors. Based on these results, we recommend concentrations of 100 µg mL^−1^ kanamycin, 25 µg mL^−1^ spectinomycin, 50 µg mL^−1^ gentamicin, 1.25 µg mL^−1^ erythromycin, and 5 µg mL^−1^ chloramphenicol for selection using the respective AbR cassettes. **(C)** Growth analysis of five putative neutral site mutants transformed with a spectinomycin resistance cassette (SpR) and grown in MAD medium as described in the Materials and Methods. Lowercase letters indicating significant difference (P < 0.05) are shown, as determined by ANOVA followed by Tukey’s honestly significant difference tests. Error bars show the mean ±SEM of three biological replicates. Abbreviations: Cm, chloramphenicol; DF, Down Flank; Em, erythromycin; Gm, gentamicin; Km, kanamycin; Sp, spectinomycin; UF, Up Flank.

Neutral sites are genomic loci that allow for stable integration of heterologous genes into the genome with no or minimal phenotypic impact. Several neutral sites have been identified in model cyanobacterial species, such as PCC 6803 and PCC 7002 (Pinto et al., 2015; Ruffing et al., 2016; Vogel et al, 2017). However, no neutral sites have yet been characterised in PCC 11901, specifically at the high densities achievable in this strain. We identified five putative neutral site loci in PCC 11901 based on previously identified neutral integration sites in other cyanobacteria and analysis of the PCC 11901 genome. *desB* (FEK_04840) encodes for a putative fatty acid desaturase, which in PCC 7002 is involved in modulating membrane fluidity at temperatures below 22⁰C but does not impact growth at 30⁰C (Sakamoto et al., 1997; Ruffing et al., 2016). *glgA1* (FEK_14880) encodes for one of two putative glycogen synthase isoforms in PCC 11901, which has previously been characterised as a neutral site in PCC 6803 and PCC 11801 (Sengupta et al., 2020; Mittermair et al., 2021). The loci for *mrr* (FEK30_09380) and *aquI* (FEK30_10065) encode for a putative Type IV restriction endonuclease and a Type II site-specific deoxyribonuclease, respectively, and were selected based on the hypothesis that several endonuclease genes may be redundant for immunity and not essential for growth. Studies in other cyanobacterial strains, such as *Thermosynechococcus elongatus* BP-1 and PCC 6803, have also reported improved transformation efficiencies in nuclease mutants (Kufryk et al., 2002; Iwai et al., 2004). Lastly, we identified an intergenic region of 185 bp between two convergent predicted open reading frames encoding hypothetical proteins (FEK30_11550 and FEK30_11555), which we hypothesised would not contain important regulatory elements. We called this locus neutral site 1 (NS1). Such regions have previously been used successfully as neutral sites in PCC 6803 and other bacterial chassis strains, including *E. coli, Bacillus subtilis* and *Pseudomonas putida* (Bernhards et al., 2022).

To establish and validate the concentrations at which selection can be performed, we chose the putative neutral site *desB*, previously tested in PCC 11901 (Mills et al., 2022), and introduced antibiotic resistance cassettes (AbRs) for each antibiotic via natural transformation (**Figure 1B**). Following three days of growth, we observed hundreds of transformant colonies for each AbR at concentrations of 100 µg mL^−1^ kanamycin, 25 µg mL^−1^ spectinomycin, 50 µg mL^−1^ gentamicin, 1.25 µg mL^−1^ erythromycin, and 5 µg mL^−1^ chloramphenicol, respectively. Based on the robust susceptibility of PCC 11901 to spectinomycin and previous success with using spectinomycin for selection in model strains (Vasudevan et al., 2019), we next assembled integrative vectors carrying a spectinomycin resistance cassette (SpR) flanked by homologous regions for each of the five target putative neutral sites to facilitate homologous recombination (HR), and transformed these into wild-type PCC 11901 to produce the mutants Δ*mrr*::SpR, Δ*aquI*::SpR, Δ*desB*::SpR, Δ*glgA1*::SpR, and Δ*NS1*::SpR (**Figure 1C**). Remarkably, we observed full segregation for each of the five mutants following a single re-streak from the transformation plates containing 25 µg mL^−1^ spectinomycin (**Supplementary Figure S2**). The segregated mutants were then subjected to a comparative growth analysis to assess the suitability of each putative neutral site. We observed that *mrr* and *aquI* were the best performing neutral sites, with Δ*mrr*::SpR and Δ*aquI*::SpR reaching optical densities similar to wild-type (i.e. OD_750_ > 100). In contrast, Δ*desB*::SpR and Δ*NS1*::SpR grew similarly to wild-type up to OD_750_ ∼50, but then growth rates declined, suggesting that these neutral sites should only be used at lower cell densities. The growth rate of Δ*glgA1*::SpR declined from OD_750_ ∼20, indicating that *glgA2* is not able to compensate for the loss of *glgA1* as observed in other strains (Sengupta et al., 2020; Mittermair et al., 2021). Notably, all five mutants grew to higher densities compared to wild-type PCC 6803 and PCC 7002. Together, our results show that endonuclease encoding genes are promising targets for identifying additional robust neutral sites in PCC 11901.

### PCC 11901 is amenable to conjugation

Delivery of heterologous DNA into genetically tractable cyanobacteria can also be achieved through conjugal transfer or triparental mating using an *E. coli* ‘helper’ strain with appropriate transmissible plasmids to enable conjugation of mobilizable plasmid vectors, such as RSF1010-based vectors (Gale et al., 2019). As genome integration and segregation are not required, conjugation has proved to be a powerful tool for rapidly testing genetic parts and libraries in several cyanobacterial strains (Bishé et al., 2019; Puzorjov et al., 2021). We sought to determine whether PCC 11901 is amenable to conjugation by introducing the empty RSF1010-based CyanoGate acceptor vectors pPMQAK1-T and pPMQAK1-T-eYFP, previously assembled by Vasudevan et al., (2019), which both contain a kanamycin resistance cassette (KmR) (**Figure 2A**). We initially obtained colonies on selective media containing kanamycin (**Figure 2B**), but subsequently found that a portion of re-streaked colonies exhibited a yellowing or chlorotic phenotype and were unable to survive successive rounds of streaking (**Figure 2C, D**). We hypothesised that the initial colonies were likely false-positives due to the native kanamycin resistance of PCC 11901 (**Figure 1A, Supplementary Figure S1**). Thus, we constructed new acceptor vectors pPMQSK1-1 and pPMQSK1-T (for level 1 and level T in CyanoGate) that carried SpR, and assembled pPMQSK1-1 vectors carrying either an eYFP (pPMQSK1-1-eYFP) or *Francisella novicida* derived Cas12a (also known as *Fn*Cpf1) expression cassette (pPMQSK1-1-Cas12a). Following conjugal transfer of pPMQSK1-1-based vectors into PCC 11901 we observed no chlorotic phenotypes in any transconjugant strains after re-streaking (**Figure 2D**), demonstrating that spectinomycin allows for robust selection of transconjugants.

**Figure 2.**
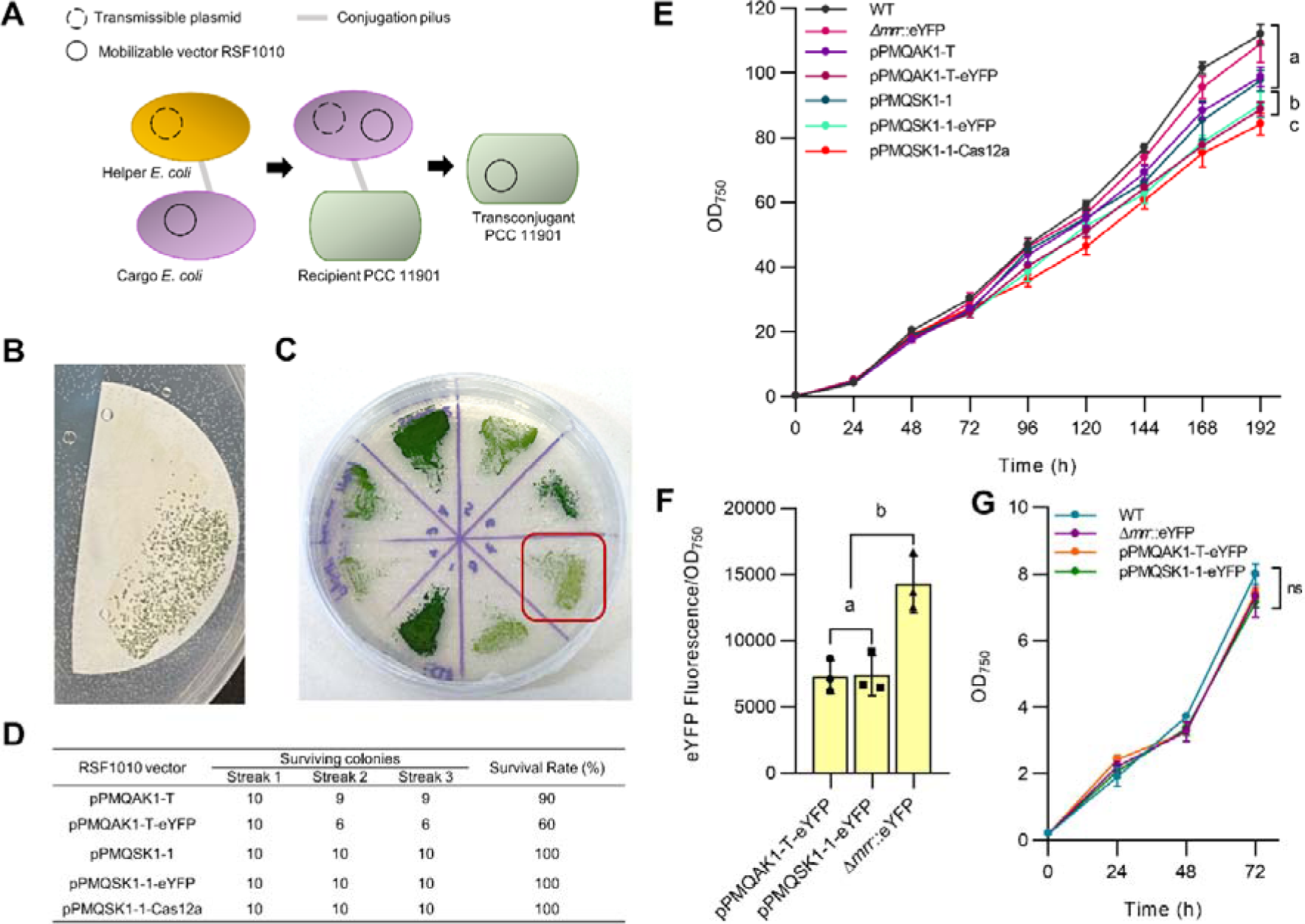
Transconjugation in PCC 11901. **(A)** Illustration showing conjugal transfer of a self-replicating RSF1010-based vector from an *E. coli* ‘cargo strain’ into PCC 11901. A transmissible helper plasmid is transferred from an *E. coli* helper strain to the cargo *E. coli* strain, which in turn aids the transfer of the mobilizable RSF1010 vector into the recipient PCC 11901. This series of steps is facilitated by the formation of conjugation pili where genetic material is transferred. **(B)** Representative image of PCC 11901 colony growth on membrane filters following selection on AD7 agar supplemented with 100 µg mL^−1^ kanamycin **(C)** pPMQAK1-T-based conjugants showed a typical dark green phenotype or a pale phenotype (red box) after streaking colonies from membranes filters onto agar media containing kanamycin. Pale colonies did not survive re-streaking. **(D)** Survival rates of transconjugants harbouring RSF1010-based vectors selected for with kanamycin (pPMQAK1-T and pPMQAK1-T-eYFP) or spectinomycin (pPMQSK1-1, pPMQSK1-1-eYFP and pPMQSK1-1-Cas12a). **(E)** Growth comparison of transconjugant and transformant strains. **(F)** Fluorescence of transconjugant and transformant strains expressing eYFP measured at 24 h. **(G)** Growth comparison for transformant and transconjugant strains expressing eYFP. For (E), (F) and (G) lowercase letters indicating significant difference (P < 0.05) are shown, as determined by ANOVA followed by Tukey’s honestly significant difference tests. Error bars show the mean ±SEM of three biological replicates.

We next performed a comparative growth analysis to explore the potential impact of RSF1010-based vectors on high density growth in PCC 11901. We included a stably integrated Δ*mrr*::eYFP mutant for comparison between transconjugants and transformants (pC1.493, **Supplementary Table S2**). The growth of transconjugants with empty acceptor vectors pPMQAK1-T or pPMQSK1-1 were not significantly different to wild-type PCC 11901 and Δ*mrr*::eYFP (**Figure 2E**). However, growth was reduced in transconjugants with pPMQAK1-T-eYFP, pPMQSK1-1-eYFP or pPMQSK1-1-Cas12a, suggesting that the expression of genes from RSF1010-based vectors could represent a metabolic burden for PCC 11901, at least at high cell densities (Meyer, 2009). A comparison of eYFP fluorescence levels between Δ*mrr*::eYFP and the transconjugants with pPMQAK1-T-eYFP or pPMQSK1-1-eYFP showed that eYFP expression was 50 ± 1% lower in both transconjugants compared to the stable integration mutant (**Figure 2F-G**). Overall, our results show for the first time that conjugation is feasible in PCC 11901. However, for this species the speed of transformation and segregation for genome integration appears more rapid than using conjugation. Furthermore, protein expression levels in stable mutants exceeded transconjugants with RSF1010-based vectors (at least for eYFP), suggesting that stable integration may be the favoured engineering choice in PCC 11901. Nevertheless, the capacity to maintain self-replicating vectors in PCC 11901 could still greatly facilitate the characterisation of genetic parts, an important aspect of strain engineering, and facilitate transient gene expression.

### Characterisation of constitutive promoters and transcriptional terminators in PCC 11901

Constitutive promoters provide a stable level of gene expression and are essential parts in the engineering toolbox of any chassis strain. Thus, we characterised a suite of 12 constitutive promoters derived from Vasudevan et al. (2019) to provide a promoter library with varying strengths for use in engineering PCC 11901. Included were constitutive promoters from PCC 6803 (P*_cpc560_* and P*_psbA2L_*) and synthetic promoters (P*_J23119_*, P*_J23115_*, P*_J23113_*, P*_J23111_*, P*_J23110_*, P*_J23103_*, P*_J23101_*, P*_V02_*, P*_V07_* and P*_trc10_*). Promoter strengths were assessed as expression cassettes driving eYFP through stable genomic integration at the *mrr* neutral site, or on the self-replicating RSF1010-based vector pPMQAK1-T. We were successful in assembling and characterising all 12 constitutive promoters following conjugation using pPMQAK1-T (**Figure 3A**). We were also able to generate and characterise 10 segregated integrative transformants at the *mrr* neutral site (**Figure 3B**). Unfortunately, we were not able to generate transformant colonies for P*_J23119_* and P*_trc10_* despite multiple attempts. Overall, the three strongest constitutive promoters were P*_cpc560_*, P*_J23119_*, P*_psbA2L_*, which is consistent with findings in other cyanobacterial species (Li et al., 2018; Vasudevan et al., 2019; Sengupta et al., 2020). The PCC 6803 *cpc* operon promoter (P*_cpc560_*) was used to drive high expression levels of heterologous genes in several model species, including PCC 6803 and PCC 7002. The P*_psbA2L_* promoter drives expression of the photosystem II reaction centre subunit D1. The high light levels used to grow PCC 11901 may support the strong levels of expression seen with P*_psbA2L_*, similar to previous work in PCC 6803 and UTEX 2973 under high light conditions (Lindberg et al., 2010; Sakurai et al., 2012; Li et al., 2018). Although the observed trend in promoter strength showed a strong correlation (R^2^ = 0.95) between those integrated at *mrr* and on pPMQAK1-T (**Figure 3C**), we found that genomic integration resulted in a 32% mean increase in eYFP expression for each promoter over those from RSF1010-based vectors. Previous work in PCC 6803 has observed contrasting results, with promoters on RSF1010-based vectors achieving a 3-fold higher level of expression compared to neutral site integrations (Ng et al., 2015). The latter finding was attributed to an increased plasmid copy-number for the RSF1010-based vector compared to the native chromosome. Thus, in PCC 11901 the copy number ratio may be lower for RSF1010-based vectors relative to the native chromosomes. The chromosomal copy number for PCC 11901 remains unclear and should be a focus for further work.

**Figure 3.**
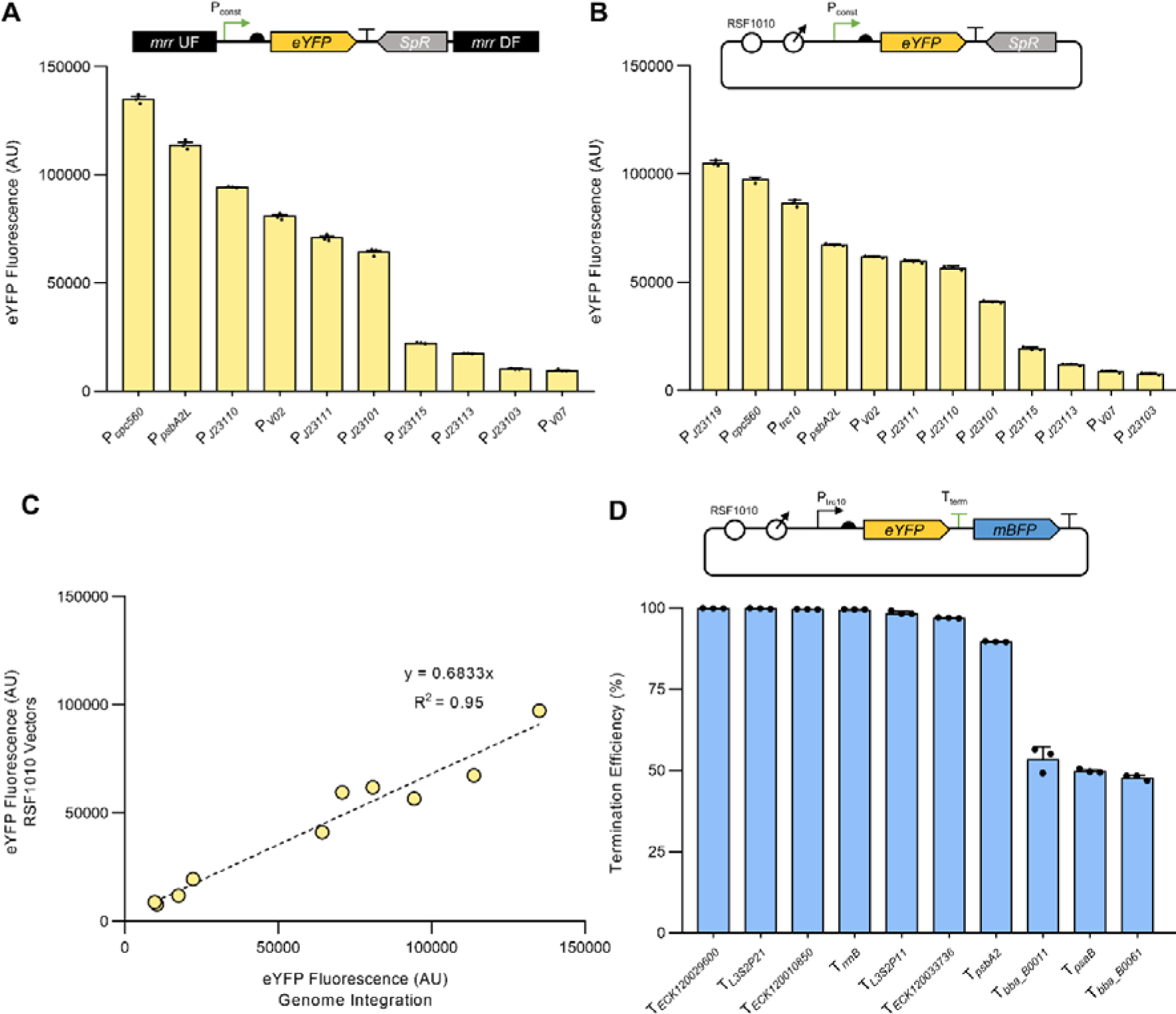
Characterisation of constitutive promoters and transcriptional terminators in PCC 11901. **(A)** Expression levels of eYFP driven by constitutive promoters integrated into the *mrr* neutral site, or **(B)** on the self-replicating pPMQAK1-T vector. **(C)** Correlation analysis of the expression levels of constitutive promoters integrated into the *mrr* neutral site or on a pPMQAK1-T vector. **(D)** Termination efficiency (TE) values for terminator sequences calculated as in Gale et al. (2021). Error bars represent the mean ±SEM of three biological replicates, each calculated from 10,000 individual cells. Abbreviations: P_const_, constitutive promoter; T_term_, transcriptional terminator.

We next characterised the efficiencies of a suite of 10 transcriptional terminators in PCC 11901 based on those previously characterised in PCC 6803, UTEX 2973 and *E. coli* by Gale et al. (2021). Cyanobacteria rely on the mechanism of rho-independent transcription termination or intrinsic termination, which is defined by the formation of a hairpin loop on the terminator sequence that leads to dissociation of the RNA polymerase and release of the mRNA transcript (Wilson and von Hippel, 1995). Our suite of intrinsic terminators included six native terminators from *E. coli* (T*_ECK120010850_*, T*_ECK120033736_*, T*_ECK120029600_*, T*_bba_B0011_*, T*_bba_B0061_*, and T*_rrnB_*), two synthetic terminators derived from *E. coli* sequences (T*_L3S2P21_* and T*_L3S2P11_*), and two native terminators from PCC 6803 from photosystem II subunit D1 (T*_psbA2_*) and photosystem I subunit B (T*_psaB_*) (Chen et al., 2013; Liu & Pakrasi, 2018). Of these, six had termination efficiency (TE) values of >95%, with highest and lowest TE values observed for T*_ECK120029600_* (99.8%) and T*_bba_B0061_* (47.8%), respectively (**Figure 3D**). Our results were relatively consistent with those from PCC 6803 and UTEX 2973, supporting our previous observation that these transcriptional terminators perform similarly across different cyanobacterial chassis (Gale et al., 2021).

### Inducible gene expression systems in PCC 11901

Chemically inducible promoters can be used to modulate gene expression in response to an external stimulus and are powerful tools for fundamental research and advanced engineering approaches (e.g. gene circuit assembly) (Meyer et al., 2019). We first tested the L-rhamnose inducible promoter P*_rhaBAD_* and its cognate transcription factor, RhaS, which has previously been characterised in PCC 6803 (Kelly et al., 2018; Behle et al., 2020). To evaluate this system in PCC 11901, we initially investigated if L-rhamnose impacted the growth of wild-type PCC 11901 over a 72 h growth period. We found that PCC 11901 growth was not affected by concentrations of up to 20 mM L-rhamnose, which was twice the highest concentration tested by Behle et al. (2020), suggesting that L-rhamnose is not toxic to PCC 11901 (**Supplementary Figure S3A**). We then constructed an RSF1010-based reporter system RhaS/P*_rhaBAD_*-*eYFP*-T*_rrnB_*in the pPMQAK1-T vector, using the medium strength constitutive promoter P_J*23101*_ to drive RhaS expression and eYFP as the fluorescent reporter (**Figure 4A**). Following successful conjugal transfer into PCC 11901, cultures were induced with increasing concentrations of L-rhamnose (0-20 mM) and eYFP fluorescence was measured after 24 h (**Figure 4B**). In the uninduced state (0 mM L-rhamnose), a low level of eYFP fluorescence was detected (305 ± 9 AU, 6.4% of the maximum expression level), demonstrating that the P*_rhaBAD_* promoter was leaky. Upon induction, we found that the promoter achieved maximum expression (4,776 ± 185 AU) with 10 mM L-rhamnose, giving a 15-fold dynamic range. Leaky expression with the P*_rhaBAD_*promoter has been reported previously in PCC 6803 (Liu et al., 2020). Those authors successfully increased the control of expression by replacing the promoter ribosome binding site with a theophylline riboswitch, although their system required two small molecules for induction.

**Figure 4.**
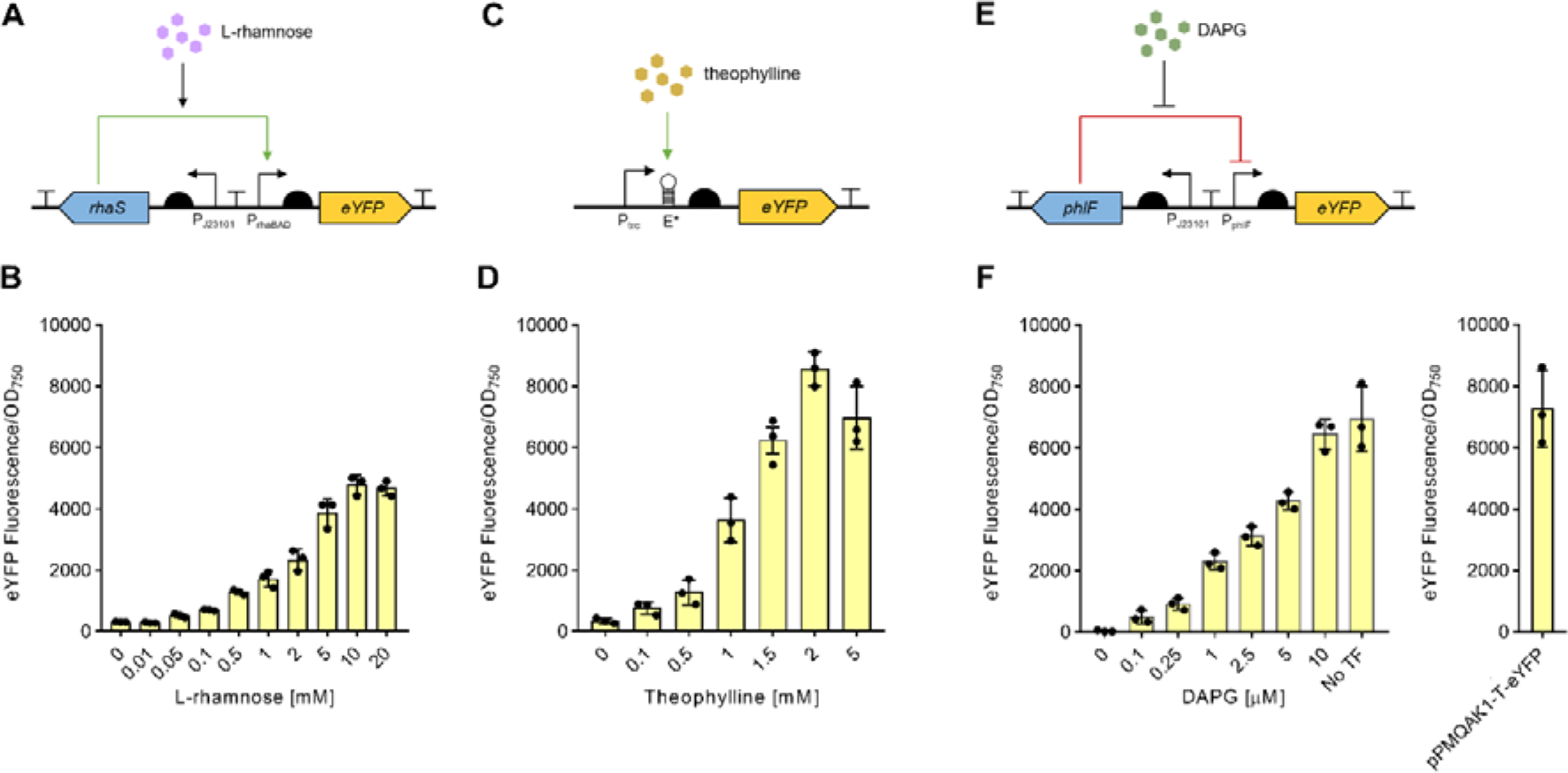
Evaluation of three inducible promoter systems in PCC 11901. Overview of the genetic components and dose-response expression levels of eYFP 24 h after induction with increasing concentrations of their respective substrate for the **(A, B)** L-rhamnose-inducible promoter RhaS/P*_rhaBAD_* fluorescent reporter system, **(C, D)** theophylline-inducible promoter P*_trc_*_E*_ fluorescent reporter system and **(E, F)** DAPG-inducible PhlF/P*_phlF_*fluorescent reporter system. The eYFP expression level for the constitutive P*_cpc560_* promoter (pPMQAK1-T-eYFP, Figure 2F) is included for comparison. Error bars show the mean ±SEM of three biological replicates.

RNA riboswitches are structured noncoding RNA domains that can regulate gene expression and exert translational control in bacteria by binding small molecules (Kavita and Breaker, 2023). We tested the engineered theophylline-inducible riboswitch E* fused to the *trc* promoter (P*_trc_*) first demonstrated in PCC 7942 (Nakahira et al., 2013). We observed that growth of wild-type PCC 11901 was not affected at concentrations up to 2 mM theophylline but was reduced at 5 mM (**Supplementary Figure S3B**). This is consistent with reports in other cyanobacteria of the negative impact of theophylline on growth at concentrations above 2 mM (Ma et al., 2014). We then assembled an RSF1010-based reporter harbouring P*_trc_*_E*\**_-*eYFP*-T*_rrnB_* in the pPMQAK1-T vector for conjugal transfer into PCC1901 (**Figure 4C**). In the absence of theophylline, an eYFP fluorescence signal above background levels was still detected in conjugant strains (336 ± 163.38 AU, 4.8% of the maximum expression level), indicating that P*_trc_*_E*_ was leaky in PCC 11901, similar to P*_rhaBAD_* (**Figure 4D**). Maximum eYFP expression (8,555 ± 315 AU) was achieved with 2 mM theophylline (giving a 25-fold dynamic range), but expression levels decreased at 5 mM theophylline, likely due to growth inhibition. The observed leaky expression with P*_trc_*_E*_ was consistent with reports in several other cyanobacterial species (Taton et al., 2017; Chi et al., 2019).

We sought to identify an inducible system for PCC 11901 that has tighter regulation than the P*_rhaBAD_* and P*_trc_*_E*_ promoters, so we next evaluated the 2,4-diacetylphloroglucinol (DAPG)-inducible promoter P*_phlF_* and its cognate transcription factor PhlF, which has tight regulation in *E. coli* but has not yet been characterised in cyanobacteria (Meyer et al., 2019). In contrast to the RhaS/P*_rhaBAD_* system, the PhlF transcription factor functions as a repressor to the P*_phlF_* promoter, which undergoes a conformational change upon binding to DAPG and releases the promoter leading to transcription, similar to the TetR family of repressors (Abbas et al., 2002). We assembled two pPMQAK1-T vectors with i) a no TF control harbouring the expression cassette PhlF/P*_phlF_*-*eYFP*-T*_rrnB_*to test the functionality of the P*_phlF_* promoter in the absence of the PhlF repressor transcription factor and DAPG, and ii) the reporter system *phlF*/P*_phlF_*-*eYFP*-T*_rrnB_*, where the medium strength constitutive promoter PJ*_23101_* was used to drive PhlF expression (**Figure 4E**). Following conjugation into PCC 11901, we observed robust levels of YFP expression for the P*_phlF_* promoter in the no TF control (94% of P*_cpc560_*) (**Figure 2F** and **4F**). For the reporter system, increasing levels of eYFP fluorescence was observed with increasing DAPG concentrations, with maximum expression (6,440 ± 279 AU) achieved upon induction with 10 µM DAPG, similar to that for the no TF control. We observed near background levels for eYFP fluorescence (28 ± 27 AU) in the uninduced state (0 µM DAPG), indicating very tight repression of P*_phlF_*by PhlF. We found that DAPG had no impact on the growth of PCC 11901 up to 10 µM, but that growth rates were reduced at 25 µM DAPG, indicating partial toxicity at higher DAPG concentrations (**Supplementary Figure S3C**). The latter result was not unexpected, as DAPG has been shown to have broad spectrum activity against bacteria and fungi, particularly those pathogenic to plants (Keel, 1992; Julian et al., 2021). To the best of our knowledge this is the first report of the PhlF/P*_phlF_*system being successfully utilised in a cyanobacterial strain. Overall, it provided a wide, 228-fold dynamic range of induction in PCC 11901 and showed tight repression in the absence of DAPG.

### Gene repression using CRISPRi

CRISPRi is now well established as a powerful tool to explore gene function and pathways in a variety of model cyanobacterial species (Liu et al., 2020; Yao et al., 2020; Dallo et al., 2023). However, the number of characterised induction systems for CRISPRi remains relatively low for cyanobacteria. Many publications utilise the TetR-based anhydrotetracycline (aTc)-inducible promoters (Huang and Lindblad, 2013), but partial leakiness (i.e. transcriptional repression in the absence of the inducer molecule) and the degree of repression of the available systems can limit effective application, depending on the strain used. Furthermore, aTc is photosensitive and degrades rapidly in UV or blue light (Zess et al., 2016), which we believed would be problematic given the light intensities used to culture PCC 11901 for high density growth and exposure to sunlight for outdoor growth.

To alleviate the potential challenge of aTc instability under high light, we sought to develop a robust and tightly regulated inducible dCas9 CRISPRi approach in PCC 11901 by testing our three inducible promoter systems. Using the CyanoGate system, we assembled the dCas9 expression cassettes *rhaS*/P*_rhaBAD_*-*dCas9*-T*_rrnB_*, P*_trc_*_E*_-*dCas9*-T*_rrnB_* and *phlF*/P*_phlF_*-*dCas9*-T*_rrnB_* in level 1 position 1 (L1P1). In level 1 position 2 (L1P2), we assembled four different single-guide RNA (sgRNA) expression cassettes (P*_trc10_*__TSS_-sgRNA-sgRNA scaffold) targeting eYFP at four different sequence locations (**Figure 5A**) (Vasudevan et al., 2019). As a control we also assembled a constitutive expression cassette with a low strength synthetic promoter (P*_J23113_-dCas9*-T*_rrnB_*), as previous work has indicated that strong expression of large proteins, such as dCas9, can produce a metabolic burden that negatively impacts growth (Depardieu and Bikard, 2020). L1P1 and L1P2 vectors were then assembled together into the level T acceptor pUC19-T pCAT.015 vector for subsequent integration into the *desB* neutral site of the Δ*mrr*::eYFP strain.

**Figure 5.**
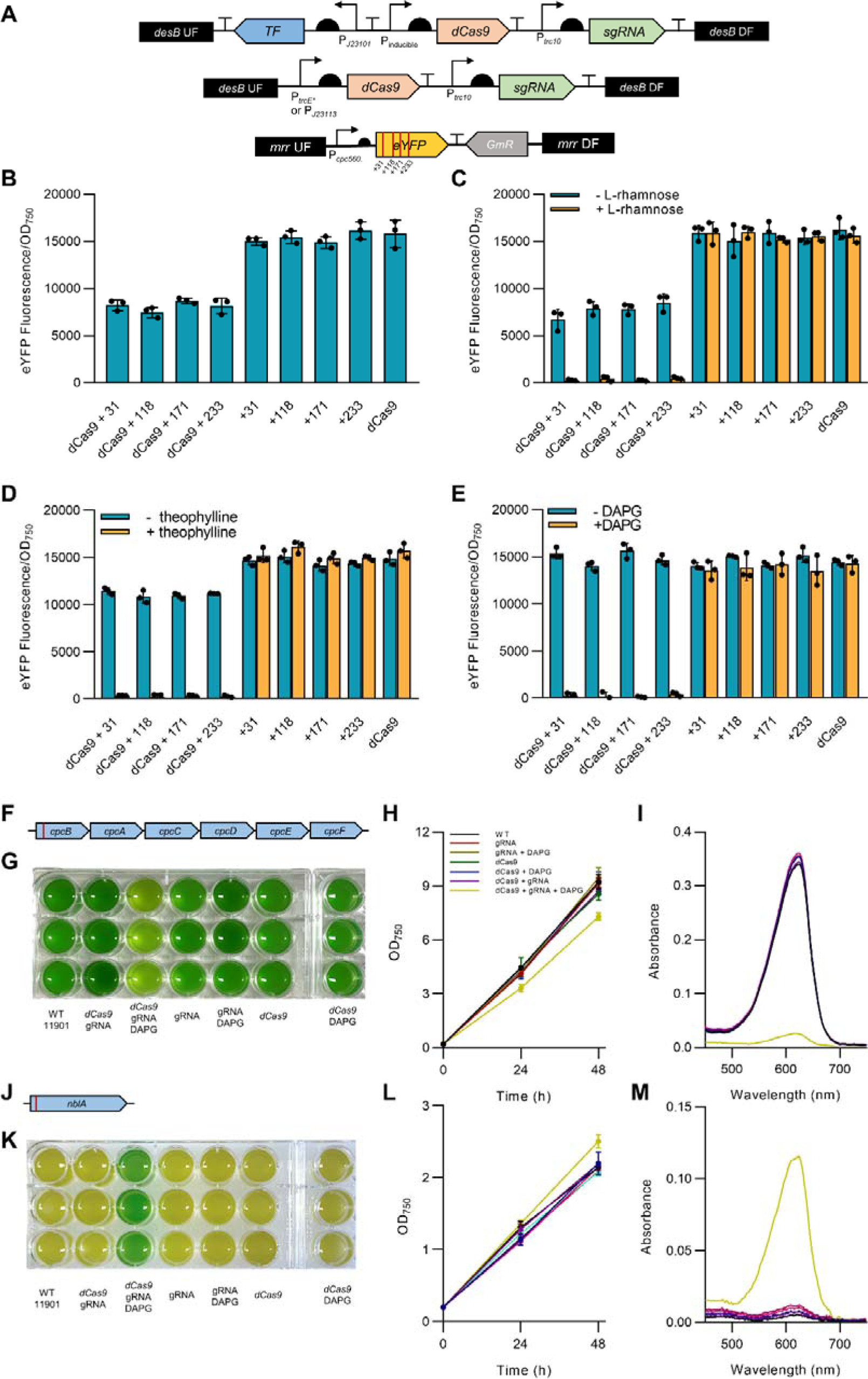
Inducible CRISPR interference (CRISPRi) for conditional knockdown of gene expression in PCC 11901. **(A)** Overview of the approaches used to test dCas9 functionality by targeting an eYFP expression cassette integrated into the *mrr* neutral site (bottom schematic) using sgRNAs targeting four different sites neighbouring a dCas9 protospacer adjacent motif sequence 5’-NGG-3’ in the eYFP open reading frame (Vasudevan et al., 2019). **(B-E)** eYFP fluorescence of plasmid vectors carrying sgRNAs with and without a constitutively expressed (P*_J23113_*) or inducible (RhaS/P*_rhaBAD_*, PhlF/P*_phlF_* and P*_trc_*_E*_) dCas9 and a strain carrying no sgRNA as a control. **(F)** Illustration of the *cpc* operon and the sgRNA target site (red bar, see **Supplementary Table S3** for sequence) in *cpcB* (c-phycocyanin subunit β, FEK30_15275). **(G)** The *cpcB* CRISPRi strains with and without inducer after 24 h of growth in MAD medium. **(H)** Growth of the *cpcB* CRISPRi strains with and without inducer over 48 h. **(I)** Absorbance spectra of PBS extracts from strains in (G). **(J)** Illustration of the sgRNA target site in *nblA* (nonbleaching A, FEK30_13550). **(K)** The *nblA* CRISPRi strains with and without inducer after 24 h of growth in MAD medium without nitrogen. **(L)** Growth of the *nblA* CRISPRi strains with and without inducer over 48 h. **(M)** Absorbance spectra of PBS extracts from strains in (K). Error bars show the mean ±SEM of three biological replicates.

We initially tested the functionality of dCas9 in PCC 11901 using the weak constitutive promoter P*_J23113_*. Following transformation and segregation, we found that only strains carrying both dCas9 and an sgRNA showed a reduction in eYFP fluorescence, which was similar for all four sgRNAs used (52 ± 3.9%) (**Figure 5B**). No significant impacts on growth were observed between strains (**Supplementary Figure S4A**). Although this demonstrated that CRISPRi-dCas9 was functional in PCC 11901, the reductions in eYFP fluorescence were unimpressive, which we attributed to the low strength of the promoter P*_J23113._* Our results are in line with those in PCC 7002 and *E. coli* where weaker promoters driving dCas9 expression resulted in lower repression of the target gene (Gordon et al., 2016; Fontana et al., 2018).

We then tested the three inducible CRISPRi-dCas9 systems by inducing transformed cultures with 10 mM L-rhamnose, 2 mM theophylline, or 10 µM DAPG, respectively, and measured the change in eYFP fluorescence. Leakiness of dCas9 expression was investigated by measuring eYFP fluorescence in the absence of the respective inducer molecules. For the RhaS/P*_rhaBAD_* CRISPRi-dCas9 system we observed a 45-58% reduction in eYFP fluorescence in the absence of L-rhamnose (**Figure 5C**), indicative of the P*_rhaBAD_* promoter leakiness observed previously (**Figure 4B**). P*_rhaBAD_* appeared able to constitutively express dCas9 at similar levels to that of the weak constitutive promoter P*_J23113_*. Overall, our results were consistent with the leakiness reported for P*_rhaBAD_* when used to express ddCas12a in PCC 6803 (Liu et al., 2020). Induction with L-rhamnose resulted in a 98 ± 1% decrease in eYFP fluorescence for all sgRNAs in the presence of dCas9. The leakiness of the theophylline-inducible P*_trc_*_E*\**_ was also apparent when used to regulate dCas9, showing a 27% reduction in eYFP fluorescence in the absence of theophylline (**Figure 5D**). Induction of dCas9 expression with theophylline resulted in a similar robust decrease in eYFP fluorescence (98 ± 1%). In contrast, cultures transformed with the DAPG-inducible PhlF/P*_phlF_*CRISPRi-dCas9 system showed no reductions in eYFP fluorescence in the absence of DAPG, while the addition of DAPG resulted in an average 97 ± 1% decrease in eYFP fluorescence for cultures with sgRNAs and dCas9 (**Figure 5E**).

Growth analyses showed that expressing dCas9 under the control of the RhaS/P*_rhaBAD_*, P*_trcE*_* or PhlF/P*_phlF_* system with their respective inducer molecules did not impair growth (**Supplementary Figure S4B-D**). Overall, our results show that the PhlF/P*_phlF_* system tightly regulated dCas9 expression with no measurable leakiness, while induction with DAPG provided robust transcriptional repression. Notably, similar levels of repression were observed for all four sgRNAs for each promoter tested. This was unexpected, as previous work in Vasudevan et al. (2019) showed different levels of repression for the same four sgRNAs in PCC 6803, and may indicate that PCC 11901 is more amenable to CRISPRi repression.

We next tested the PhlF/P*_phlF_* CRISPRi-dCas9 system on two endogenous gene targets in PCC 11901 to characterise its capacity for affecting native metabolism. Our gene targets were *cpcB* and *nblA*, which are involved in the synthesis and degradation of phycobiliproteins, respectively, and have been targeted previously in CRISPR-based studies in PCC 7002 and UTEX 2973 (Gordon et al., 2016; Ungerer and Pakrasi, 2016; Wendt et al., 2016). The *cpc* operon is involved in the synthesis of the phycocyanin rods of the phycobilisome complex (**Figure 5F**), an assemblage of proteins that bind to photosystem I or II and increase the efficiency of light harvesting (Puzorjov and McCormick, 2020; Domínguez-Martín et al., 2022). We designed an 18 bp sgRNA targeting *cpcB* (**Supplementary Table S3**), the first gene of the *cpc* operon, and checked to ensure that no off-target sites were present in the genome of PCC 11901 (Bae et al., 2014). Following induction with DAPG, we found that the CRISPRi strain expressing dCas9 and the sgRNA targeting *cpcB* had turned a yellow-green colour after 24 h of induction, similar to the olive colour observed in phycobilisome-deficient PCC 6803 mutants (**Figure 5G**) (Lea-Smith et al., 2014; Vasudevan et al., 2019), and its growth was reduced compared to the controls (**Figure 5H**). A significant decrease in the absorption peak at 625 nm was also observed in phycobilisome extracts from the latter strain (**Figure 5I**), demonstrating a robust reduction in phycobilisome abundance. SDS-PAGE analysis of phycobilisome extracts showed a decrease in both *cpc* and *apc* peptides in the induced (but not uninduced) CRISPRi strain, indicating that CRISPRi repression of *cpcB* resulted in a reduction in overall phycobilisome abundance in PCC 11901 (**Supplementary Figure S5A**). The apparent co-repression of allophycocyanin was unexpected, as previous *cpc* mutants in PCC 6803 showed no reduction of allophycocyanin (Lea-Smith et al., 2014; Liberton et al., 2017). However, previous attempts to knockout *cpcB* in PCC 7002 (the closest relative species to PCC 11901) failed, suggesting that *cpcB* is essential in PCC 7002 (Gordon et al., 2016). Our results indicate that phycocyanin and allophycocyanin may be co-regulated in PCC 11901, which may also be the case in PCC 7002. Measurement of chlorophyll content in the CRISPRi strains showed no significant differences (**Supplementary Figure S5B**), suggesting that the abundance of other components of the light reactions were not affected by the repression of *cpcB*. Finally, we tested if removal of DAPG could derepress *cpcB* and restore the phycobilisome pool. Washing the cells in fresh media to remove DAPG resulted in increased growth after 24 h and restoration of phycobilisome abundance to wild-type levels after 48 h (**Supplementary Figure S5C** and **S5D**). Thus, our results indicate that ∼24 h is sufficient for the turnover and degradation of the dCas9 pool in PCC 11901.

We next targeted the *nblA* gene, which plays a key role in phycobilisome degradation under nitrogen deficient conditions (**Figure 5J, Supplementary Table S3**) (Collier and Grossman, 1994). *nblA* is a common target to test CRISPR-Cas functionality in cyanobacteria as disruption of *nblA* leads to an easily detectable phenotype under nitrogen-limiting conditions; wild-type strains grown in media lacking nitrate exhibit chlorosis or bleaching characteristic of phycobilisome degradation, while *nblA* mutants show a non-bleaching phenotype and remain blue-green (Wendt et al., 2016; Baldanta et al., 2022; Cengic et al., 2022). Following induction with DAPG in nitrogen-depleted medium, we found that the CRISPRi strain expressing dCas9 and the sgRNA targeting *nblA* remained blue-green, whereas all the controls turned chlorotic (**Figure 5K**). All cultures grew very slowly due to the lack of nitrogen (**Figure 5L**), which is not unexpected as nitrogen plays a crucial role in the fast-growing phenotype of PCC 11901 (Włodarczyk et al., 2020). Chlorophyll content was reduced overall (**Supplementary Figure S5E**), indicating that all cultures were stressed. Nevertheless, a strong absorption peak at 625 nm was maintained in phycobilisome extracts from the functional CRISPRi strain, while all other controls showed a much-reduced peak indicative of phycobilisome degradation (**Figure 5M**).

Overall, the PhlF/P*_phlF_* inducible CRISPRi system shows tight regulation of dCas9 activity and robust repression of heterologous and native gene expression. Furthermore, we have shown that removal of the inducer DAPG can reversibly attenuate gene repression and related physiological effects rapidly in PCC 11901. This system should provide a powerful tool for studying gene function in PCC 11901 and enable metabolic engineering approaches to exploit the yield potential of PCC 11901 (Li et al., 2016; Yao et al., 2020; Miao et al., 2023).

### Markerless gene editing using CRISPR-Cas12a and a ‘double HR’ approach

The generation of markerless mutants has key advantages for biotechnology as strains can be repeatedly genetically manipulated and the absence of genes encoding antibiotic resistance proteins avoids the possibility of antibiotic resistant organisms being released into the environment (Lea-Smith et al., 2016). However, recent efforts to use the common negative selection markers *sacB* and *codA* to generate markerless mutants proved unsuccessful in PCC 11901 (Mills et al., 2022). To overcome this, we attempted to use a CRE-lox recombination system approach recently demonstrated in PCC 7002 (Jones et al., 2021). This system first involved generation of marked mutants via insertion of an AbR flanked by two LoxP sites, lox71 (5’-ATAACTTCGTATAATGTATGCTATACGAACGGTA-3’) and lox66 (5’-TACCGTTCGTATAATGTATGCTATACGAAGTTAT-3’) into a target site. A plasmid vector encoding CRE and a second AbR was then introduced into marked mutants and integrated into an essential locus (*rbcLXS* or *psbEFJL*) with CRE under control of the native promoter. As these genes are essential, the strains were maintained in a partially segregated state under antibiotic selection. Subsequent expression of CRE resulted in excision of the lox71-AbR-lox66 cassette from the genome. Growth of these mutants on plates lacking the second antibiotic resulted in loss of chromosomes containing the CRE/AbR insertion and generation of markerless mutants. We attempted to replicate this system in PCC 11901. Marked, segregated mutants targeting five different loci (*ctaDIEI*, *ctacII*, *acs*, *ldhA*, *sdhA*) were successfully generated using a gentamicin resistance cassette (GmR) or KmR flanked by lox66 and lox71 (**Supplementary Figure S6**). We then generated two plasmids that allowed recombination of CRE and SpR into the *rbcLXS* or *psbEFJL* locus. However, despite repeated attempts to transform these plasmids into each of the five markerless mutants, we were unable to obtain spectinomycin resistant colonies. Moreover, no colonies were obtained when the plasmids were introduced into wild-type cells. Our inability to generate markerless mutants in PCC 11901 using this approach suggests that CRE is either toxic or that the generation of partially segregated mutants in key essential genes is extremely challenging.

In conjunction, alternate CRISPR-Cas gene editing approaches for generating markerless mutants were trialled. These allow for more efficient engineering in cyanobacteria without the requirement for positive selection markers. As the double stranded breaks generated by Cas proteins are lethal to most microbes (including cyanobacteria) due to their general lack of a non-homologous end joining pathway (Su et al., 2016), selection is based on the uptake of supplied repair template sequences to replace the sgRNA target locus by HR (Behler et al., 2018). Due to the documented cytotoxicity of Cas9 in cyanobacteria (Li et al., 2016; Racharaks et al., 2021), much CRISPR-Cas work to date has focused on reportedly non-toxic Cas12a (*Fn*Cpf1), which has led to several successful examples of gene editing in a variety of cyanobacteria (Lin et al., 2021; Baldanta et al., 2022). Recently, an RNA riboswitch-inducible dCas9 system was demonstrated in PCC 6803 that could overcome the toxicity of Cas9 (Cengic et al., 2022).

Here, we sought to build a novel CyanoGate-compatible inducible CRISPR-Cas12a system that allows for the generation of markerless mutants and multiplex editing at high efficiencies. Our system is comprised of two vectors i) an RSF1010-based (pPMQSK1-1) self-replicating ‘editing vector’ carrying Cas12a (pC1.509, **Supplementary Table S1**), and ii) a pUC19-T ‘hybrid suicide vector’ designed to deliver an sgRNA expression cassette into the editing vector for stable expression and a repair template with homology to the sgRNA target locus on the genome by two separate HR events, which we called the ‘double HR’ approach (**Figure 6A**). We chose the DAPG-inducible PhlF/P*_phlF_* promoter system to drive expression of Cas12a, which we have previously shown to be tightly regulated in PCC 11901 (**Figure 4** and **Figure 5**), and conjugated the editing vector into PCC 11901 to generate an ‘editing strain’. To assist with assembling the hybrid suicide vector, we constructed the CyanoGate-compatible sgRNA assembly acceptor vector pCA0.421 based on the CRATES system from Liao et al., (2019), which enables one-pot assembly of single or multiplexed sgRNA arrays for targeted gene editing (**Supplementary Table S1**). We hypothesised that with this approach new hybrid suicide vectors carrying one or more sgRNAs and up to six repair templates could be rapidly generated and iteratively transformed into the editing strain to generate markerless mutants with single or multiple chromosomal alterations.

**Figure 6.**
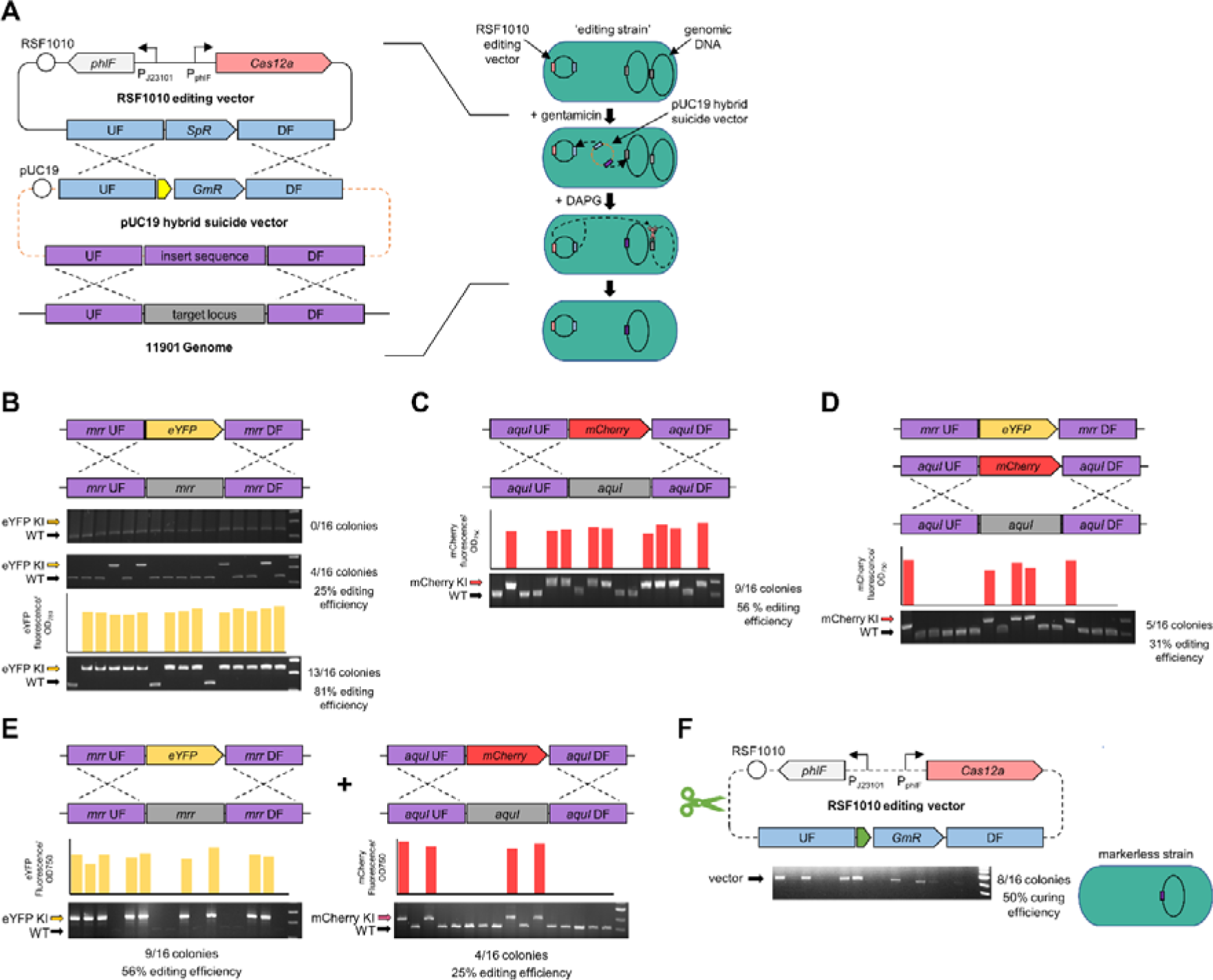
Genome editing of PCC 11901 with CRISPR-Cas12a using a double homologous recombination approach. **(A)** The editing strategy relies on an RSF1010-based editing vector (pC1.509, **Supplementary Figure S1**) and a pUC19 suicide vector. Transformation of the pUC19 suicide vector into an ‘editing strain’ carrying pC1.509 results in (1) HR with pC1.509 to deliver the sgRNA(s) and GmR (components in blue) and (2) HR with the target genomic locus to deliver a template for homology directed repair (components in purple). Subsequent DAPG-induction of Cas12a results in cleavage of the target locus (in grey), leaving only edited copies of the genome (in purple) intact (for a detailed methodology and protocols see **Supplementary Figure S7** and **S8**). **(B,C)** Demonstration of single genome editing through markerless insertion of eYFP into the *mrr* locus and mCherry into the *aquI* locus, respectively. (**D**) Iterative genome editing through markerless insertion of mCherry into the *aquI* locus of the Δ*mrr*::eYFP strain. (**E**) Multiplexed genome editing through simultaneous insertion of eYFP and mCherry into the *mrr* and *aquI* locus, respectively. (F) Demonstration of curing of the RSF1010 editing vector using a self-targeting sgRNA.

As a proof-of-concept, we used the editing strain (carrying the editing vector with SpR) to perform a markerless knock-in of an eYFP expression cassette into the *mrr* neutral site (**Figure 6B**). A hybrid suicide vector pCT.590 (**Supplementary Table S2**) was assembled and transformed into PCC 11901 in the absence or presence of DAPG, and the transformation mix was then plated onto gentamicin-supplemented agar or gentamicin- and DAPG-supplemented agar, respectively. In the absence of DAPG, no edits were observed in the colonies that grew, indicating that Cas12a expression was tightly repressed. However, with DAPG we observed that 25% of colonies showed insertions of the eYFP expression cassette and eYFP fluorescence. This demonstrated that in the Δ*mrr*::eYFP colonies i) HR had occurred between the hybrid vector and the editing vector to allow for growth on gentamicin, and ii) HR had occurred between the hybrid vector and the genome. In the remaining 75% of colonies, only i) had occurred, which indicated that HR with the editing vector was more prominent than with the genome. We confirmed that double HR had occurred with colonies expressing eYFP by PCR of the editing vector, which showed the presence of the sgRNA cassette and that SpR had been replaced by GmR (**Supplementary Figure S9**). Notably, the absence of a WT band in the Δ*mrr*::eYFP colonies indicated that these strains were fully segregated.

To further improve the efficiency of editing, we performed transformation in the absence of DAPG for 4 h, then added DAPG for 12 h before plating onto gentamicin- and DAPG-supplemented agar to induce Cas12a expression. We observed a significant increase in the efficiency of double HR (81%) (**Figure 6B**), which demonstrated the benefit of increasing the time period for genomic HR to occur prior to the induction of Cas12a expression. This approach was used in all subsequent gene editing experiments. We next assembled the hybrid suicide vector pCT.621 to test the *aquI* neutral site as a second target locus for insertion of an mCherry expression cassette (**Supplementary Table S2**). Here we observed a 56% efficiency in generating fully segregated Δ*aquI*::mCherry lines (**Figure 6C**), suggesting that the double HR approach was still robust at different genomic loci.

We next investigated if we could generate the double insertion mutant Δ*mrr*::eYFP/Δ*aquI*::mCherry through iterative or multiplex genome editing. For the former, we assembled a hybrid suicide vector pCT.622 for transformation into Δ*mrr*::eYFP, which carries an editing vector with GmR (**Supplementary Table S2**). Following transformation of Δ*mrr*::eYFP, we observed mCherry expression cassette insertions in *aquI* in 31% of colonies (**Figure 6D**). Fluorescence measurements confirmed expression of both eYFP and mCherry, and thus successful generation of Δ*mrr*::eYFP/ Δ*aquI*::mCherry double mutants through iterative gene editing. For the latter multiplex approach, we assembled the suicide vector pCT.623 carrying an sgRNA array and homology repair templates designed to insert expression cassettes for eYFP and mCherry into *mrr* and *aquI*, respectively. Following transformation of the editing strain, we screened colonies for single and double insertion mutants by PCR. We observed insertion efficiencies of 56% and 25% for the eYFP and mCherry expression cassettes, respectively, while 25% of the colonies contained both cassettes at their expected loci (**Figure 6E**). Notably, we did not find examples of an mCherry insertion in the absence of eYFP, suggesting that the first sgRNA in the array targeting the *mrr* locus may have been more efficient or is more abundantly expressed.

Overall, our results support PCC 11901 as a highly amenable strain for CRISPR-Cas12a-based editing. To our knowledge, this is the first report of iterative CRISPR-Cas12a gene editing and multiplex editing using an sgRNA array in a cyanobacterial strain. The iterative editing scheme was designed to allow as many sites to be targeted as desired, by cycling through SpR and GmR cassettes in the hybrid suicide vector, enabling more complex editing schemes to occur. The two editing schemes (iterative and multiplex) could also be further combined, allowing further flexibility in strain design. Additional efficiency improvements could be made by establishing the mechanism by which ‘escaper colonies’ survive on selective media but avoid editing, a common phenomenon observed in bacteria and cyanobacteria (Vento et al., 2019; Cengic et al., 2022). As we were able to achieve iterative gene editing, escape was not due to mutations in Cas12a and thus may have been due to point mutations in the sgRNA or the sgRNA genomic target site. A possible strategy to improve the efficiency of edits could be to reduce the expression levels of Cas12a and/or the sgRNA(s), an approach that that has been shown to increase editing efficiencies in *E. coli* and *Klebsiella* spp. (Collias et al., 2023).

Finally, we sought to cure the editing strain of the self-replicating editing vector. Various methods have been employed to cure plasmids from bacteria, with the standard approach being repeated subculture of mutants in antibiotic-free medium and screening for spontaneous plasmid loss (Bishé et al., 2019). However, RSF1010-based vectors appear to persist in cyanobacteria for long periods, even in the absence of antibiotic selection (Nagy et al., 2021; Puzorjov et al., 2022). Previous CRISPR-Cas work in *E. coli* and *P. putida* utilised a ‘self-targeting’ sgRNA for efficient removal of RSF1010-based vectors (Lauritsen et al., 2017). Here, we generated a hybrid suicide vector containing an sgRNA that targeted the editing vector pC1.509. We transformed cells in the presence of DAPG and plated the transformation culture onto agar without antibiotics. We then screened colonies for the absence of the editing vector by PCR and observed a curing efficiency of 50% (**Figure 6F**). We further verified the loss of the editing vector by patching the cured colonies onto gentamicin-supplemented agar, which resulted in no growth indicating sensitivity to the antibiotic. Thus, we demonstrated that PCC 11901 mutants generated by CRISPR-Cas12a can be cured of the editing vector to produce fully markerless mutant strains containing no scars or AbR cassettes.

## CONCLUSION

Here we have investigated the amenability of the fast-growing marine cyanobacterium PCC 11901 to engineering and assembled a comprehensive suite of tools compatible with the CyanoGate MoClo platform to enable future work in this strain. We hope that the tools described will accelerate the wider adoption of this next-generation cyanobacterial chassis strain. The fast-growing and highly productive phenotype of PCC 11901 offers much in terms of advancing our fundamental understanding of the genetic basis and regulation of these processes (Ungerer et al., 2018). Furthermore, PCC 11901 shows promise for applied work aiming to develop commercially viable green biotechnology chassis for renewable biomanufacture and biomaterials production (Goodchild-Michelman et al., 2023), sequestration of CO_2_ emissions in hard-to-abate sectors (e.g. capture of CO_2_ from point-source flue gases) (Zhang et al., 2017), and sustainable space exploration (Santomartino et al., 2023).

## MATERIALS AND METHODS

### Cyanobacterial culture conditions

*Synechococcus* sp. PCC 11901 was cultured in AD7 or MAD liquid medium (Włodarczyk et al., 2020), or on 1.5% (w/v) agar plates as described in (Włodarczyk et al., 2020). Cultures were grown in an Algaetron AG 230 incubator (Photon Systems Instruments) at 30°C, 2% (v/v) CO_2_ under continuous warm white LED light (150 µmol photons m^−2^ s^−1^) and shaking at 120 rpm. Agar plates were incubated under identical conditions, without shaking.

### Plasmid vector assembly

Level 0, 1, and T plasmid vectors were assembled using the CyanoGate MoClo kit (Vasudevan et al., 2019). Native PCC 11901 genetic parts were amplified from genomic DNA using Q5 High-Fidelity DNA Polymerase (New England Biolabs). Where necessary, native genetic parts were domesticated (i.e. sites for Type IIS restriction endonucleases BsaI and BpiI were removed) using specific primers. Alternatively, parts were synthesized as Gblocks DNA fragments (Integrated DNA Technologies) and cloned directly into an appropriate level 0 acceptor (Engler et al., 2014) (see **Supplementary Table S1** and **Supplementary Data S1** for new vectors, and **Supplementary Table S2** for all plasmids assembled in this study). Vectors were transformed into One Shot TOP10 chemically competent *Escherichia coli* (Thermo Fisher Scientific) cells as per the manufacturer’s instructions. Transformed cultures were grown at 37°C on 1.5% (w/v) LB agar or in liquid LB medium shaking at 125 rpm with appropriate antibiotic selection.

### sgRNA selection and CRISPR-Cas assemblies

sgRNAs were designed by selecting 18-22 bp sequences adjacent to the protospacer adjacent motif (PAM) sequence 5’-NGG-3’ for *Streptococcus pyogenes* dCas9 or 5’-TTTV-3’ for *Francisella novicida* Cas12a. Candidate sgRNAs were checked for potential off-target sites in the PCC 11901 genome using Cas-OFFinder (Bae et al., 2014). The sgRNAs for dCas9 were made by annealing complementary oligonucleotides carrying the required overhangs and BsaI recognition sites (**Supplementary Table S3**), and were assembled into the level 1 position 2 (L1P2) acceptor vector pICH47742 together with the P*_trc10__*_TSS_ promoter (pC0.220) and the sgRNA scaffold (pC0.122) as described in Vasudevan et al., (2019). The sgRNAs for Cas12a were also made by annealing complementary oligonucleotides carrying overhangs and BpiI recognition sites as described in Liao et al. (2019) (see **Supplementary Figure S3** and **S7**) and were assembled into the new acceptor vector pC0.421 (**Supplementary Table S1**).

### Natural transformation of PCC 11901

Purified plasmid (1 µg) was added to 1 mL of wild-type PCC 11901 culture at exponential growth phase (OD_750_ = 0.8) and incubated for 12 h at 30°C under continuous warm white LED light (150 µmol photons m^−2^ s^−1^) and shaking at 120 rpm in an Infors Multitron Pro incubator (Infors HT). The cultures were then plated onto AD7 agar plates supplemented with appropriate antibiotics (25 μg mL^−1^ spectinomycin, 50 μg mL^−1^ gentamicin, 100 μg mL^−1^ carbenicillin or 100 μg mL^−1^ kanamycin). Plates were then sealed with 3M Micropore® tape to allow for gas exchange and incubated at 30°C, 2% (v/v) CO_2_ under warm white LED light (150 µmol photons m^−2^ s^−1^). Colonies typically appeared after 2-4 days.

### Transconjugation of PCC 11901

Genetic modification by conjugal transfer was performed using an approach adapted from Gale et al. (2019). Overnight cultures of *E. coli* strain HB101 harbouring vectors pRK2013 (ATCC 37159) and pRL528 (the helper strain) and a TOP10 strain harbouring an RSF1010-derived level T vector were each washed three times with LB medium to remove antibiotics. The *E. coli* cultures were then combined (450 μL each) and incubated for 1 h at room temperature (RT). PCC 11901 cultures were grown to OD_750_ ∼1.0 and washed three times with fresh AD7 medium. The combined *E. coli* culture was added to 900 μL of PCC 11901 culture and the mixture incubated at 30°C under warm white LED light (150 µmol photons m^−2^ s^−1^) for 4 h, without shaking. The mix was centrifuged at 4,000 *g* and the cell pellet was plated onto 0.45 μm Metricel® membrane filter discs (Pall Corporation) laid on top of non-selective AD7 agar. After 24 h of incubation, the membranes were transferred to AD7 agar supplemented with appropriate antibiotics (25 μg mL^−1^ spectinomycin, or 100 μg mL^−1^ kanamycin) and incubated as above. Conjugant colonies typically appeared six days post membrane transfer.

### Comparative growth assays

Growth curve experiments were performed by inoculating a seed culture containing 30 mL of MAD medium with single colonies of cyanobacteria picked from agar plates and grown as described above to OD_750_ ∼1.0. The seed cultures were then used to prepare triplicate 15 mL starter cultures adjusted to OD_750_ ∼0.2 and aliquoted into Corning® 25 cm^2^ cell culture flasks with canted necks and vented caps (Corning). To facilitate gas exchange and prevent foaming, 0.5 μL of Antifoam 204 (Sigma Aldrich) was added to the cultures. Cultures were grown at 30°C, 2% (v/v) CO_2_, shaking at 150 rpm under 150 µmol photons m^−2^ s^−1^ for the first 24 h, which was then increased to 750 µmol photons m^−2^ s^−1^. Optical density was measured every 24 h using a WPA Biowave II UV-Vis spectrophotometer (Biochrom) for eight days.

### eYFP quantifications

Mutant PCC 11901 strains were grown in 6-well culture plates (Starlab CytoOne) and incubated in an Algaetron® AG 230 incubator under the same culturing conditions as described above. OD_750_ and eYFP fluorescence of cultures were measured using a FLUOstar OMEGA microplate reader (BMG Labtech). Fluorescence of eYFP for individual cells (10,000 cells per culture) was measured by flow cytometry using a BD® LSR II Fortessa flow cytometer (Becton Dickinson). Cells were gated using forward and side scatter, and median eYFP fluorescence was calculated from excitation/emission wavelengths 488 nm/515–545 nm (Kelly et al., 2018), and reported after 24 h of growth unless otherwise stated.

### Terminator efficiency calculations

The efficiency of terminator sequences (terminator efficiency (TE)) was calculated by assembling terminator sequences into the pDUOTK1-L1 vector as described in Gale et al., (2021).

### Measurement of chlorophyll content

Cultures were diluted to OD_750_ = 1.0 and centrifuged at 17,000 *g* for 2 min. The resulting pellet was resuspended in 100% (v/v) methanol and shaken at 2400 rpm for 1 h in the dark at RT using an IKA-VIBRAX-VXR bead beater. The homogenates were then centrifuged at 17,000 *g* for 10 min and the absorbance of the supernatant was measured at 652, 665 and 750 nm. The mean concentration of chlorophyll *a* was calculated from triplicates as described in Porra et al., (1989).

### Extraction of phycobiliproteins and analysis

Phycobiliproteins (PBS) were extracted and quantified using absorbance spectroscopy as described previously (Zavřel et al., 2018). Briefly, PCC 11901 cells were pelleted by centrifugation at 15,000 *g* for 5 min, washed in phosphate buffered saline three times and freeze-dried overnight. Dried samples were lysed with 0.5 mm glass beads (BioSpec Products) on a TissueLyser II homogeniser (Qiagen) for 15 s at RT. 1 mL of pre-cooled phosphate buffered saline at 4°C was added to each tube and mixed for 5 s on the homogeniser. Samples were then incubated on ice for 60 min to efficiently extract soluble proteins and prevent protein degradation. Following centrifugation for 5 min at 4°C and 15,000 *g*, the aqueous blue liquid layer containing PBS was transferred to sterile 1.5 mL microcentrifuge tubes and frozen for future use. Samples were measured from 550-750 nm on a Biochrom WPA Biowave II Spectrophotometer. For SDS-PAGE analysis, samples were run on a Bolt 12% Bis-Tris Plus Mini protein gel (Invitrogen) at 150 V for 1 h. A pre-stained protein standard (Proteintech) was used as a ladder. The gels were then stained with 1% (w/v) Coomassie Brilliant Blue solution (Bio-Rad) and destained with a methanol: acetic acid: distilled water (50%: 10%: 40%) solution.

## ACKNOWLEDGEMENT

We would like to thank the Frankenberg-Dinkel Lab (RPTU Kaiserslautern-Landau) for kindly providing their facilities for several experiments while hosting A.J.V.

## DATA AVAILABILITY

Plasmids in **Supplementary Table S1** are available from Addgene (https://www.addgene.org/Alistair_McCormick), Addgene IDs: 203934-203943, 205441.

## SUPPLEMENTARY DATA

**Supplementary Table S1.**
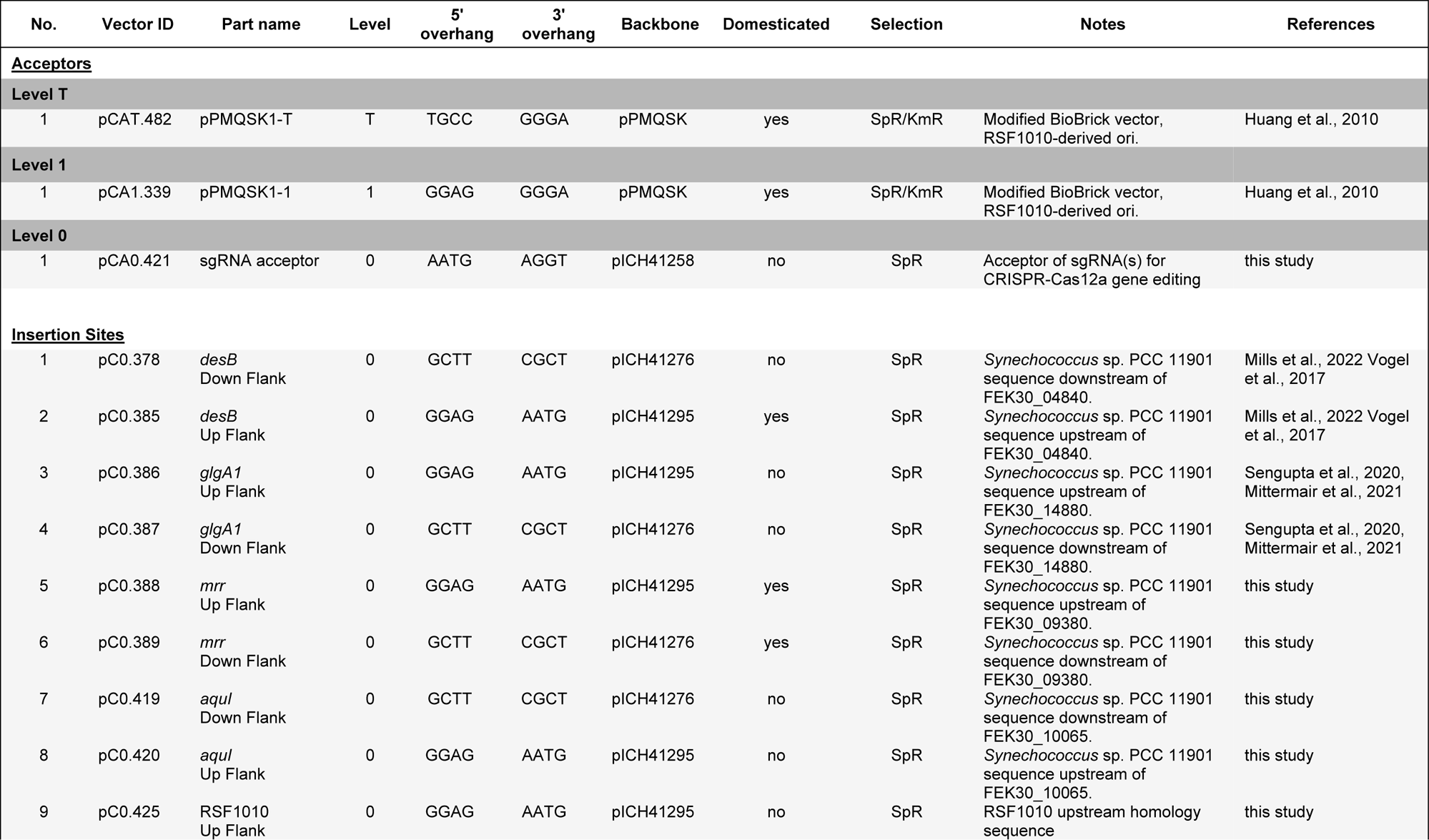
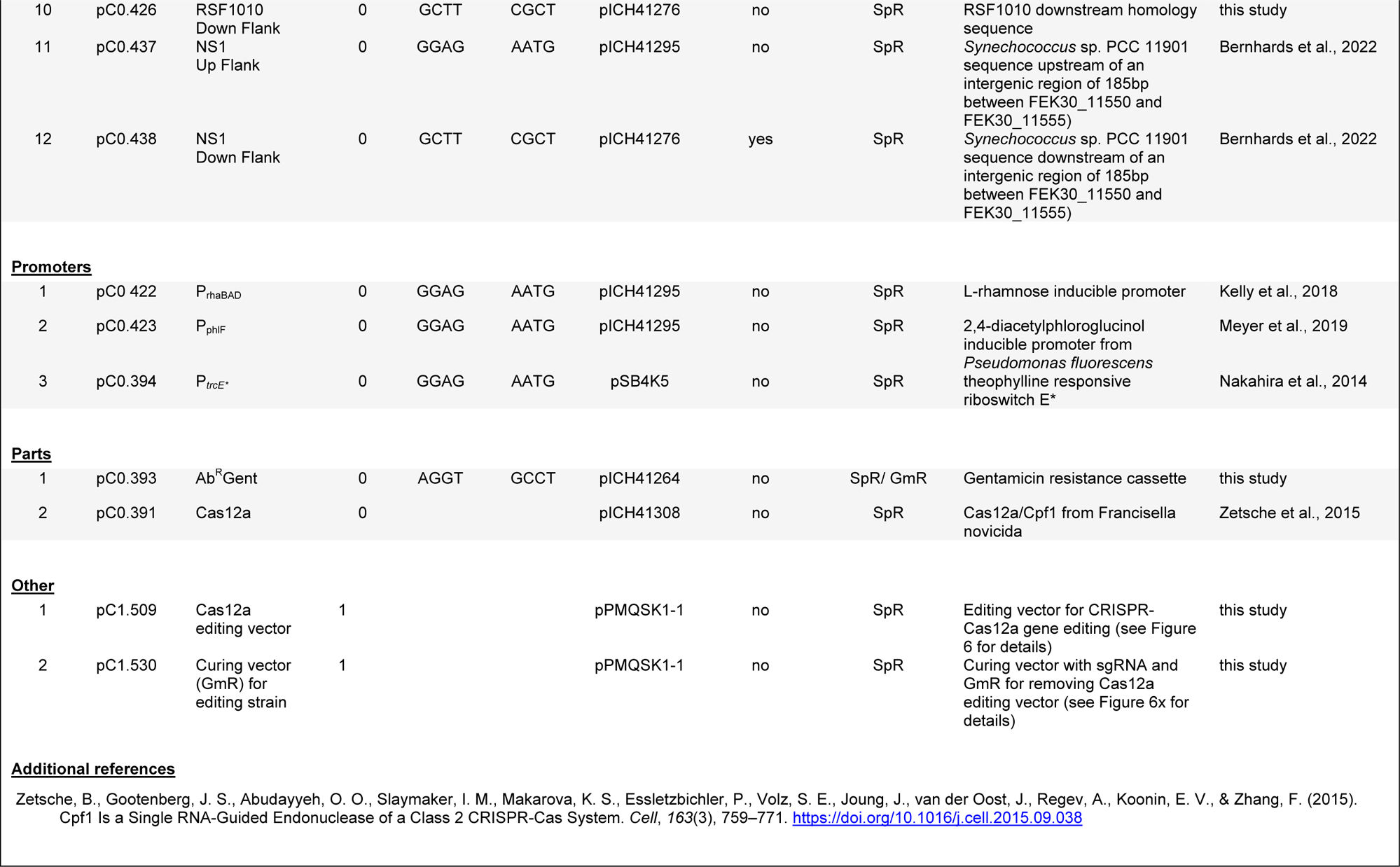
Table of all new CyanoGate-compatible parts generated in this work.

**Supplementary Table S2.** All plasmids made and used in this study. See ‘Supplementary Table S2 - Plasmids (all).xlsx’. RSF1010 vectors used for promoter and terminator characterisations in Figure 3B and 3C, respectively, have been published previously (Vasudevan et al., 2019; Gale et al., 2021).

**Supplementary Table S3.** Primer and sgRNA oligonucleotides used in this study. See ‘Supplementary Table S3 - Primer and sgRNA oligonucleotides.xlsx’.

**Supplementary Data S1.** Sequence maps (.gb files) for plasmid vectors in Supplementary Table S1. See ‘Supplementary Data S1.zip’.

**Supplementary Figure S1.**
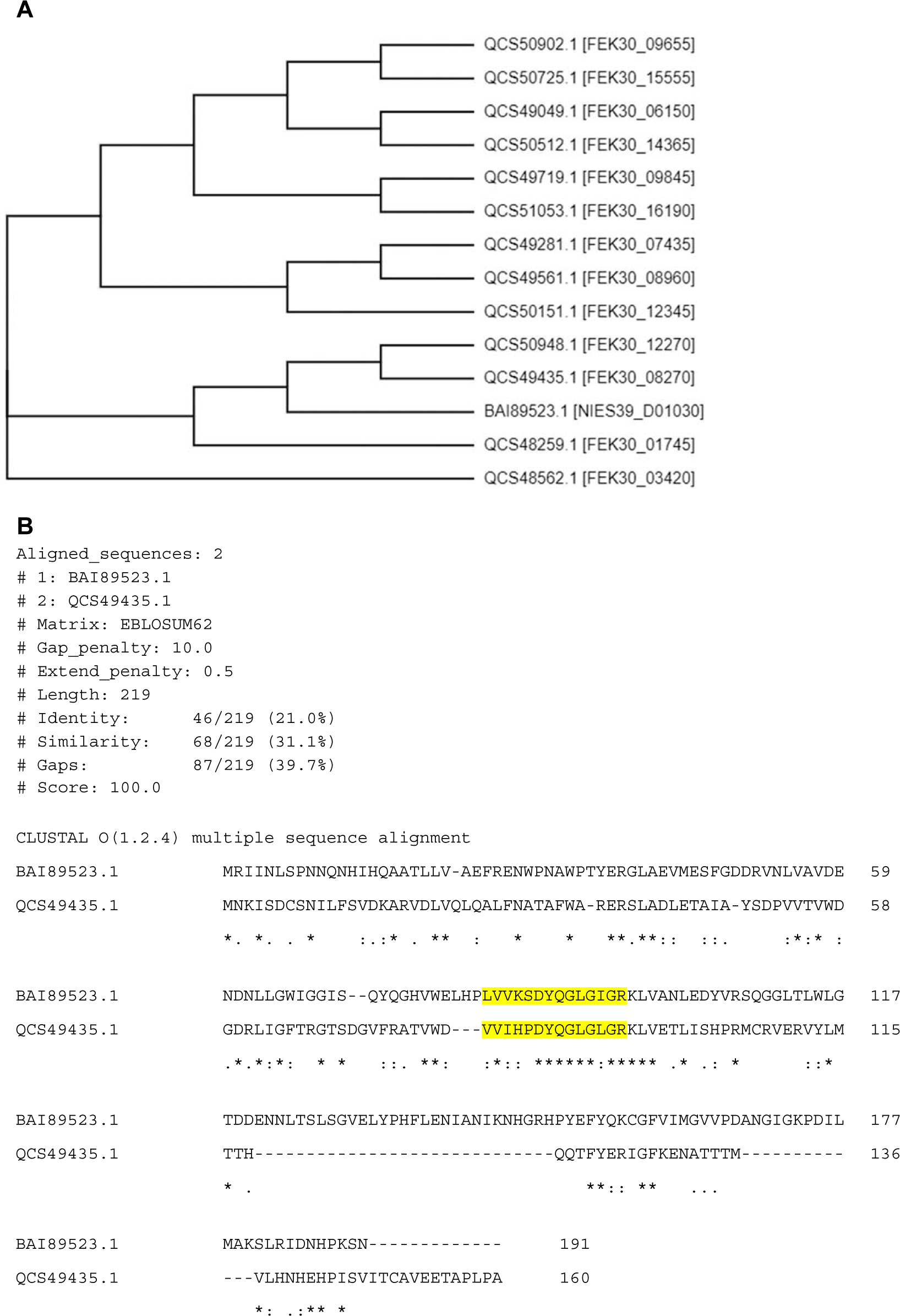
Analysis of putative Gcn5-related N-acetyltransferase (GNAT) family genes in PCC 11901. **(A)** Phylogenetic tree of 13 GNAT family proteins in PCC 11901 and the aminoglycoside acetyltransferase from *Arthrospira platensis* (BAI89523.1) (built by Clustal Omega (EMBL-EBI)). **(B)** Sequence alignment and scores between BAI89523.1 and a putative aminoglycoside acetyltransferase from PCC 11901 (QCS49435.1). The conserved coenzyme A binding pocket is highlighted in yellow. The alignment was performed in EMBOSS Needle.

**Supplementary Figure S2.**
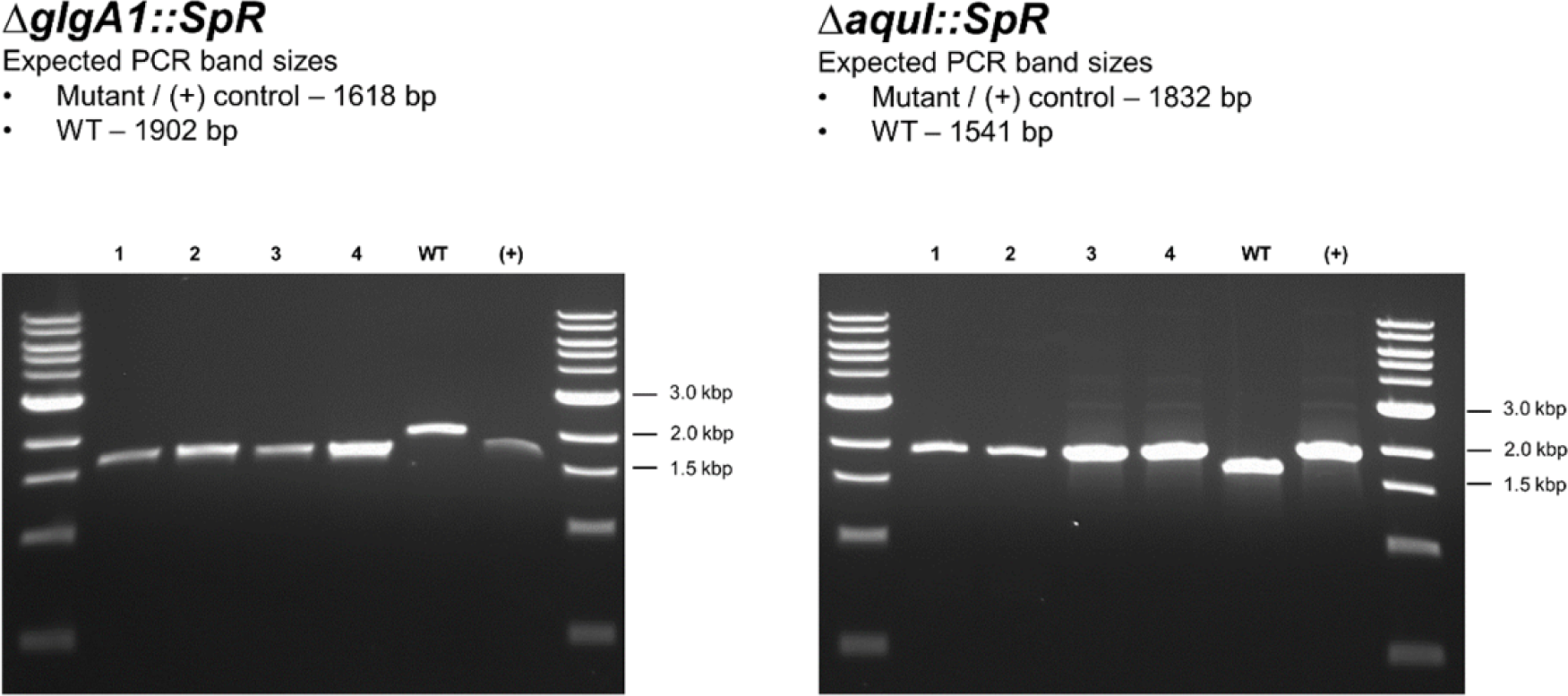
Representative PCR-based segregation analysis of PCC 11901 transformants targeting neutral sites *glgA1* and *aquI*. Electrophoresis gel profiles are shown for four independent transformant colonies after one round of re-streaking on selective agar. Full segregation for each transformant is demonstrated by the presence of amplicons showing insertion of the SpR cassette at the *glgA1* (1,618 bp) and *aquI* (1,832 bp) loci and the absence of a respective wild-type (WT) amplicon (1,902 bp for *glgA1* and 1,541 bp for *aquI*). Amplicons from each transformation plasmid (+) are included as positive controls.

**Supplementary Figure S3.**
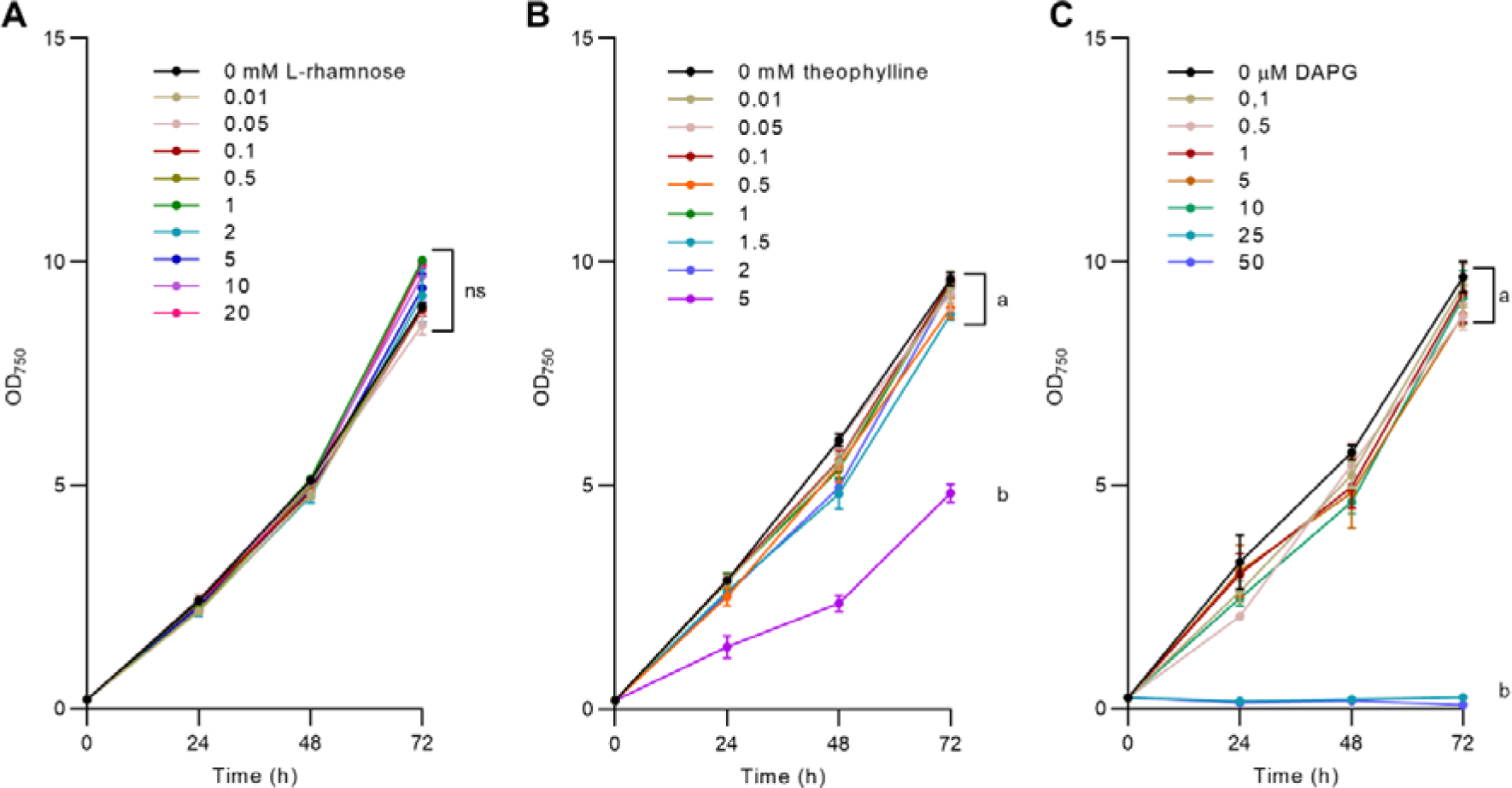
Growth of PCC 11901 with varying doses of small molecule inducers. Growth of wild-type PCC 11901 in increasing concentrations of **(A)** L-rhamnose, **(B)** theophylline, and **(C)** DAPG over 72 h. Error bars represent the mean ±SEM of three biological replicates.

**Supplementary Figure S4.**
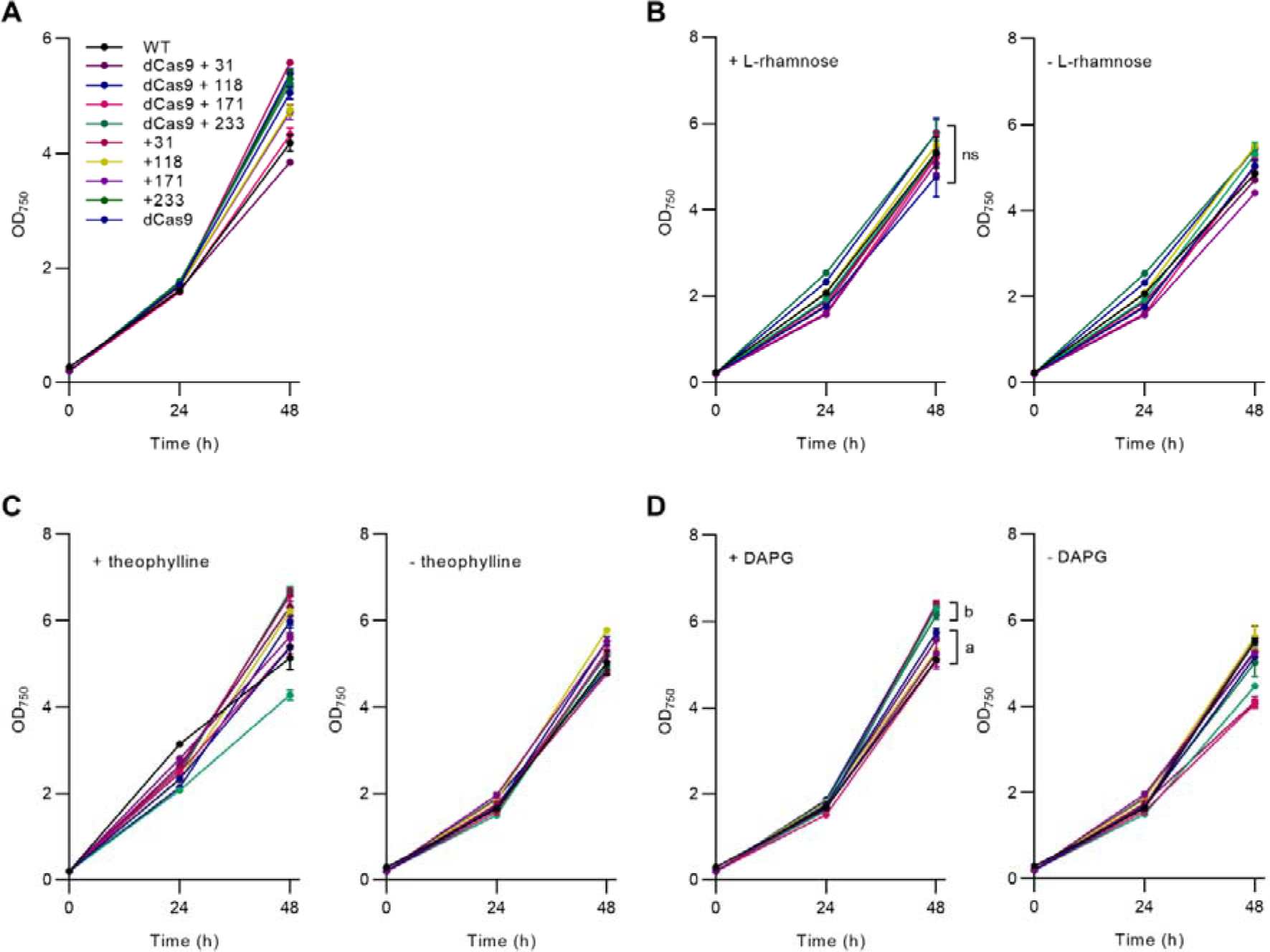
Growth analysis of CRISPRi-dCas9 strains targeting eYFP. **(A)** Growth of P_J23113_ CRISPRi strains in MAD medium. **(B)** Growth of RhaS/P*_rhaBAD_* CRISPRi strains in the presence or absence of 10 mM L-rhamnose. **(C)** Growth of P*_trcE*_*CRISPRi strains in the presence or absence of 2 mM theophylline. **(D)** Growth of PhlF/P*_phlF_* CRISPRi strains in the presence or absence of 10 µM DAPG. Error bars represent mean ±SEM of three biological replicates.

**Supplementary Figure S5.**
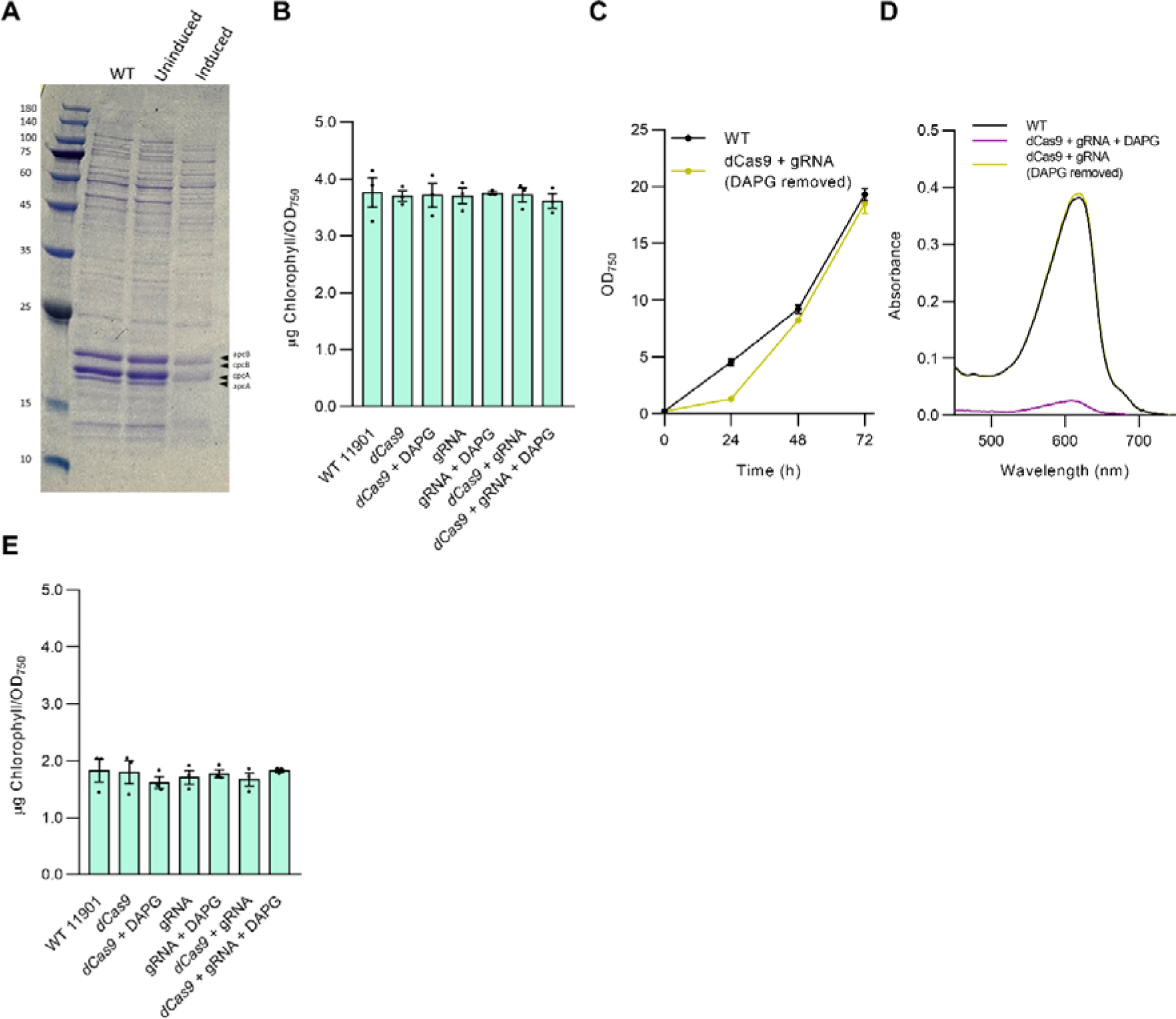
Analysis of CRISPRi-dCAs9 strains targeting *cpcB* and *nblA*. **(A)** Coomassie-stained SDS-PAGE gel of PBS extracts from the *cpcB* CRISPRi strain carrying both sgRNA and dCas9 in the absence (uninduced) or presence (induced) of 10 µM DAPG. Arrows indicate the bands for *apcB* (18.7 kdA), *cpcB* (18.1 kdA), *cpcA* (17.6 kdA) and *apcA* (17.3 kdA). **(B)** Chlorophyll contents of the *cpcB* CRISPRi strains. **(C)** Growth of the *cpcB* CRISPRi strain carrying both sgRNA and dCas9 in comparison to wild-type PCC 11901 following removal of DAPG. **(D)** Absorbance spectra of PBS extracts following removal of DAPG at 48 h. **(E)** Chlorophyll contents of *nblA* CRISPRi strains. Error bars show the mean ±SEM of three biological replicates for B, C and E.

**Supplementary Figure S6.**
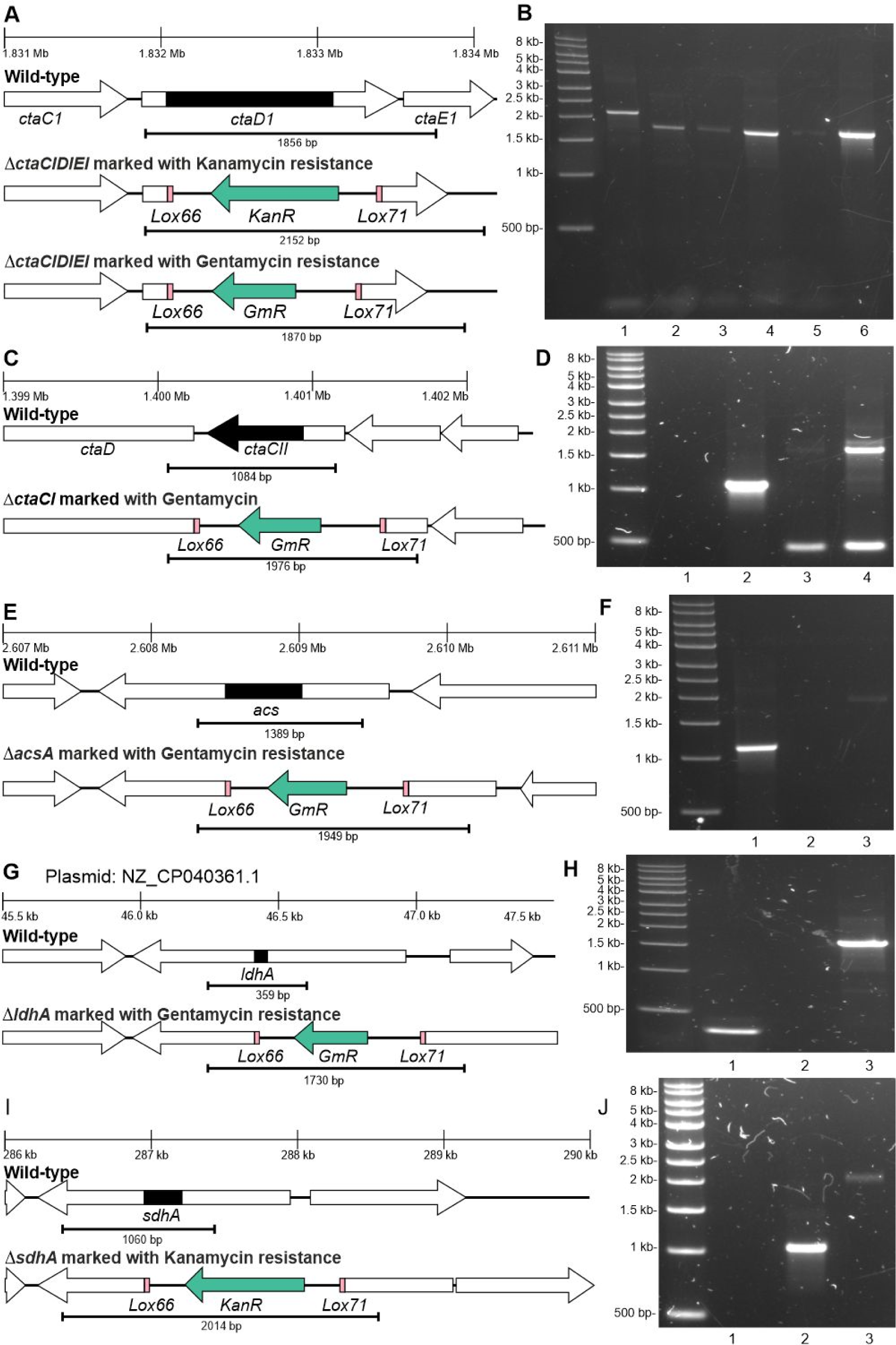
Attempted generation of markerless mutants using the CRE-Lox system. Schematic representations of locus location in the PCC 11901 genome (top) and the wild-type (middle) and marked knockout (KO, bottom) profiles expected in the **(A)** *ctaDIEI*, **(C)** *ctaCII*, **(E)** *acs*, **(G)** *ldhA* and **(I)** *sdhA* strains following amplification with primers flanking the deleted sequence. Regions deleted in the mutant strains are shaded in black. Amplification of genomic DNA in: **(B)** Δ*ctaDIEI* KmR marked KO (lane 1); Δ*ctaDIEI* GmR marked KO (lanes 2-5); WT (lane 6); **(D)** negative control (lane 1); WT (lane 2); Δ*ctaCII* GmR marked KO (lanes 3-4); **(F)** WT (lane 1); negative control (lane 2); Δ*acs* GmR marked KO (lane 3); **(H)** WT (lane 1); negative control (lane 2); Δ*ldhA* GmR marked KO (lane 3); **(J)** negative control (lane 1); WT (lane 2); Δ*sdhA* KmR marked KO (lane 3). Δ*ctaDIEI* GmR marked KOs were sequenced to confirm the correct KO profile.

**Supplementary Figure S7.**
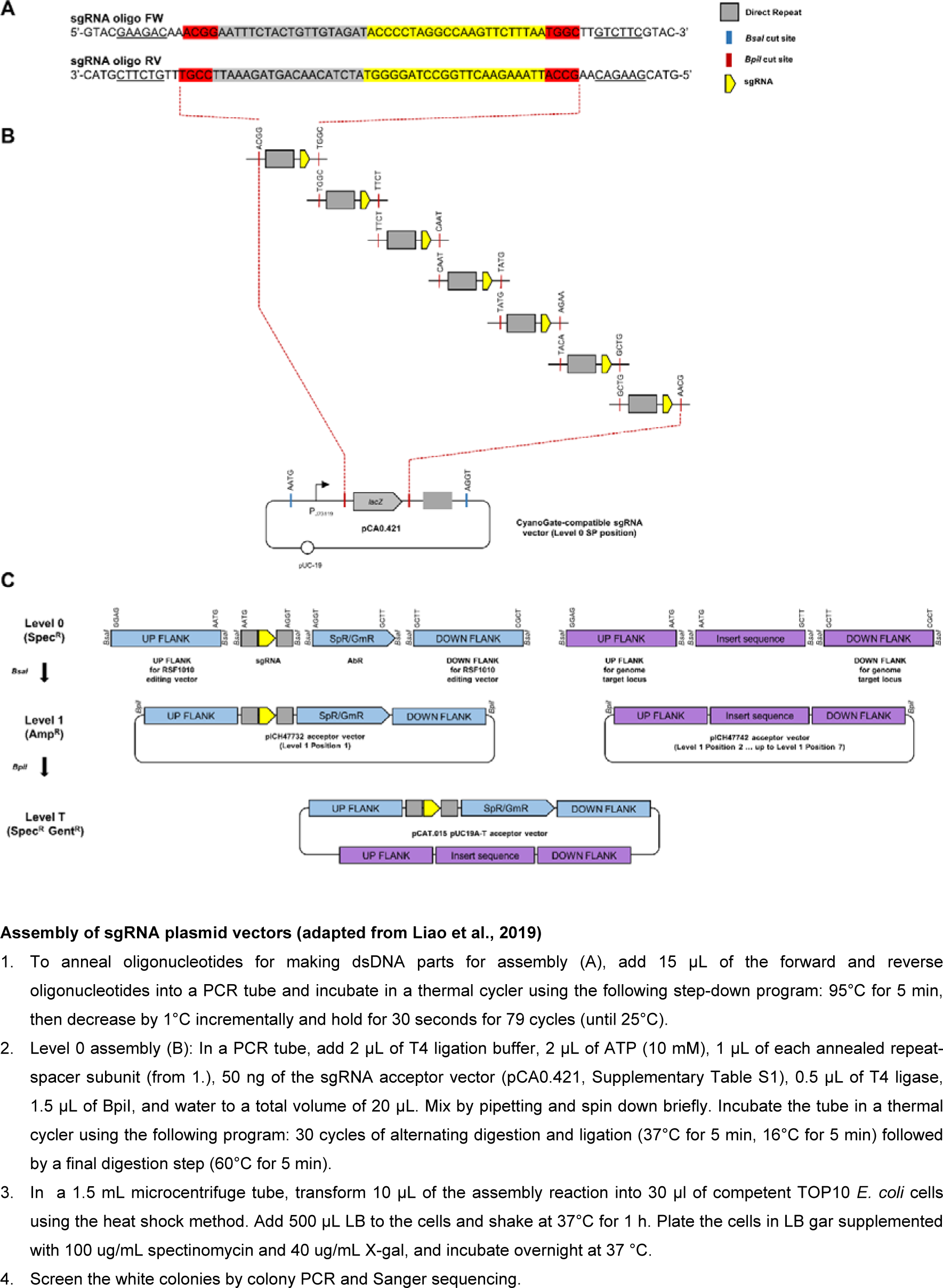
CRISPR-Cas12a double HR editing approach – sgRNA, repair template and hybrid suicide vector assembly. **(A)** sgRNA were assembled by annealing oligonucleotides incorporating the desired 18-22bp sgRNA (yellow), direct repeat (grey), unique 4 bp overhangs (red) and BpiI sites (underlined) for **(B)** assembly into the pC0.421 sgRNA acceptor vector. Overhangs match those from the CRATES system and allow for assembly of an array of between one and seven sgRNAs (Liao et al., 2019). **(C)** MoClo assembly of level 0 parts into level 1 vectors and then a level T hybrid suicide vector incorporating the sgRNA(s) in position 1 and homology repair template(s) in position 2 up to position 7 using the available Plant MoClo level 1 acceptor vectors (Engler et al. 2014). Using this system between one and six genomic loci can be targeted simultaneously. A protocol for assembly of the sgRNA plasmid vectors is included.

**Supplementary Figure S8.**
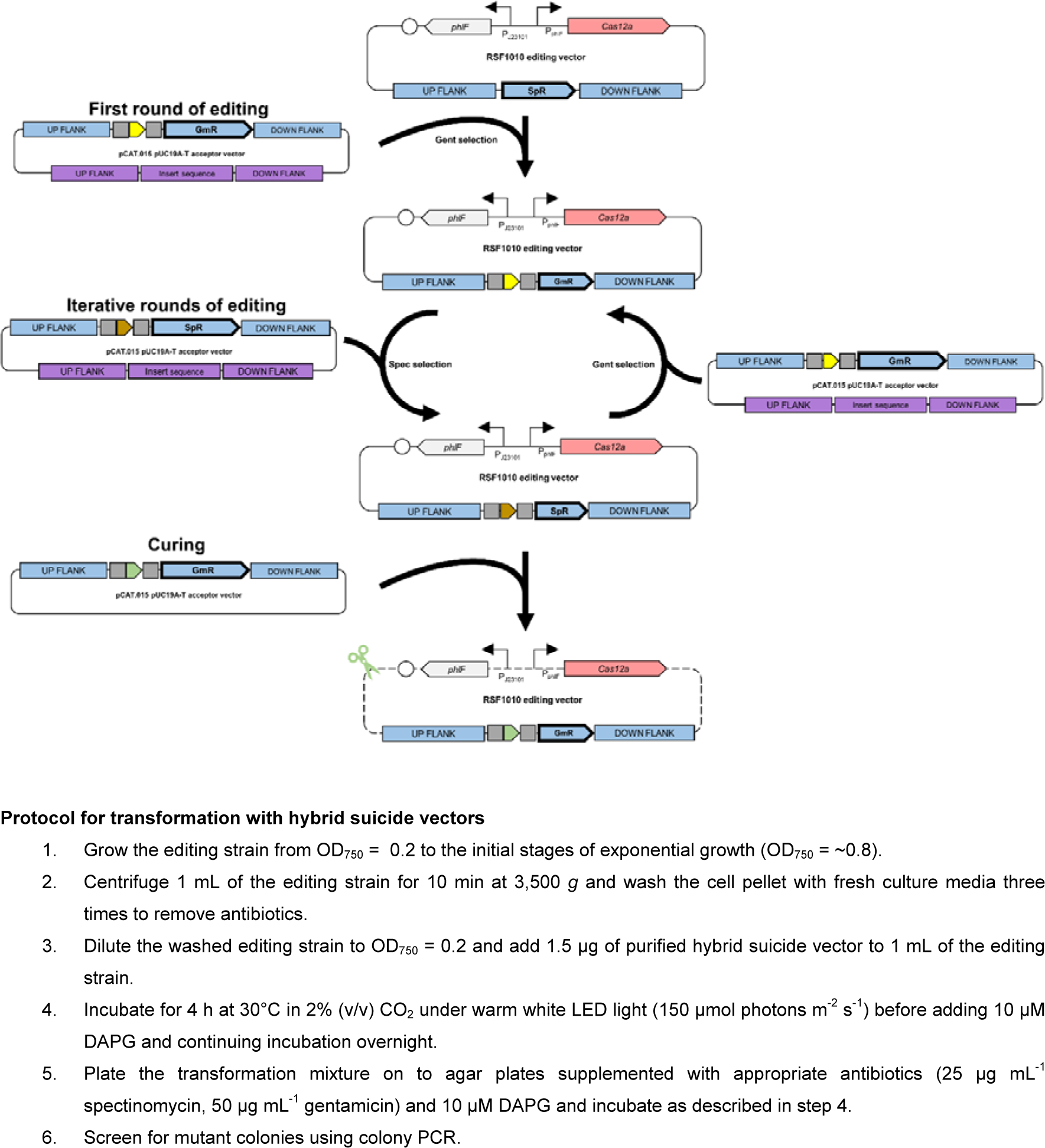
Pipeline for iterative CRISPR-Cas12a editing using the double HR approach. In the first round of editing the hybrid suicide vector recombines with the editing vector to introduce an sgRNA (or sgRNA array) and replaces the SpR with GmR for subsequent selection on gentamicin-supplemented agar plates (also see Figure 6A). Subsequent rounds of editing involved cycling between hybrid suicide vectors that exchange SpR and GmR for delivery of new sgRNA(s) to the editing vector. Curing the edited strain of the editing vector to generate a fully markerless strain is done using a hybrid suicide vector (e.g. pC1.530, **Supplementary Table S1**), which delivers a self-targeting sgRNA to the editing vector. A protocol for transformation with the hybrid suicide vector and induction with DAPG is included.

**Supplementary Figure S9.**
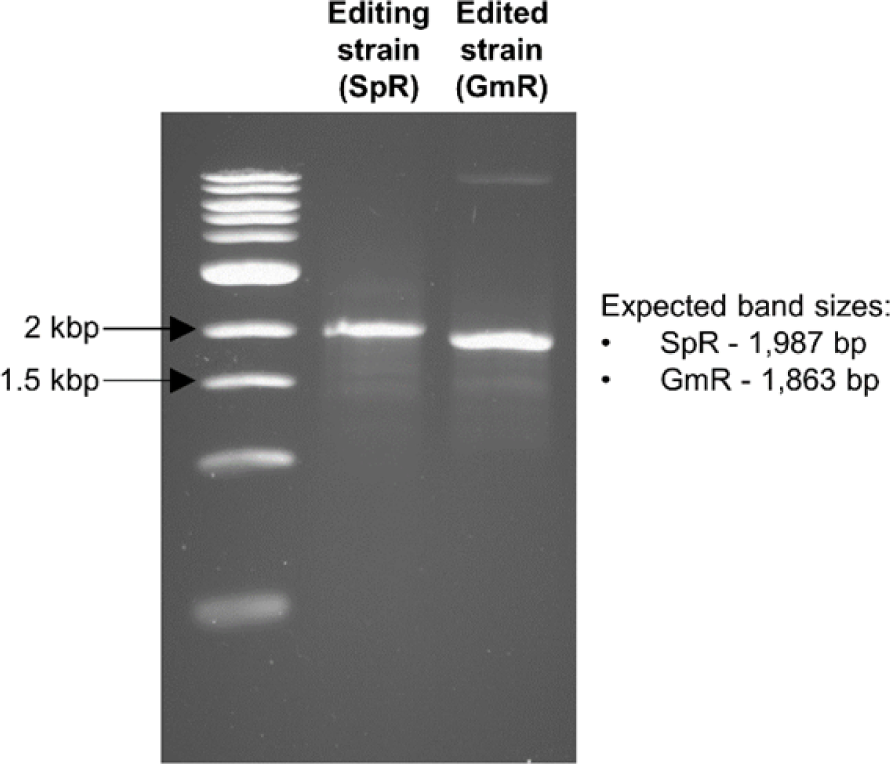
Confirmation of recombination between the hybrid suicide vector and editing vector. Example colony PCR of a transformant showing replacement of SpR by GmR

